# Monensin and Its Analogs Exhibit Activity Against Breast Cancer Stem-Like Cells in an Organoid Model

**DOI:** 10.1101/2025.07.05.663311

**Authors:** Alicja Urbaniak, Billie Heflin, Eric Siegel, Megan R. Reed, James S. Nix, Eric U. Yee, Marta Jędrzejczyk, Greta Klejborowska, Natalia Stępczyńska, Adam Huczyński, Marius B. Nagalo, Timothy C. Chambers, Steven Post, Robert L. Eoff, Melanie C. MacNicol, Amit K. Tiwari, Thomas Kelly, Alan J. Tackett, Angus M. MacNicol

## Abstract

Monensin (**MON**) is a polyether ionophore antibiotic of natural origin and is an FDA-approved drug for veterinary use. Recent studies have highlighted its potential anti-cancer activity in various *in vitro* and *in vivo* models. In this study, we evaluated the anti-breast cancer activity of **MON** and 37 synthetic analog compounds using cell monolayer and organoid models. Through a mini-ring cell viability assay, several compounds were identified that were more potent and selective against breast cancer cells compared to non-cancerous cells, surpassing the activity of parent **MON**. **MON** and these compounds induced significant DNA fragmentation, reduced cell migration, and downregulated SOX2 expression. Furthermore, **MON** and the most potent analog, compound **12**, reduced the percentage of CD44^+^/CD24^-/low^ stem-like cells and diminished cell self-renewal properties. Proteomics analyses revealed that several pathways, including extracellular matrix organization, were significantly dysregulated by **MON** and compound **12** in breast cancer cells. Among these, TIMP2, a protein associated with the suppression of tumor growth and metastasis, was identified as one of the most prominently upregulated proteins by **MON** and compound **12** in MDA-MB-231 cells. This finding was also validated in other breast cancer and melanoma cell lines. To simulate breast cancer metastasis to the brain, a human Hybrid Organoid System: Tumor in Brain Organoid (HOSTBO) model was developed. **MON** and compound **12** significantly reduced Ki-67 expression within the HOSTBOs, and compound **12** significantly downregulated SOX2 expression. Collectively, **MON** and compound **12** significantly reduced the proliferation of breast cancer stem-like cells in the organoid models, inhibited their migration, and dysregulated markers associated with stemness, demonstrating their potential as anti-metastatic agents and warranting further clinical development.

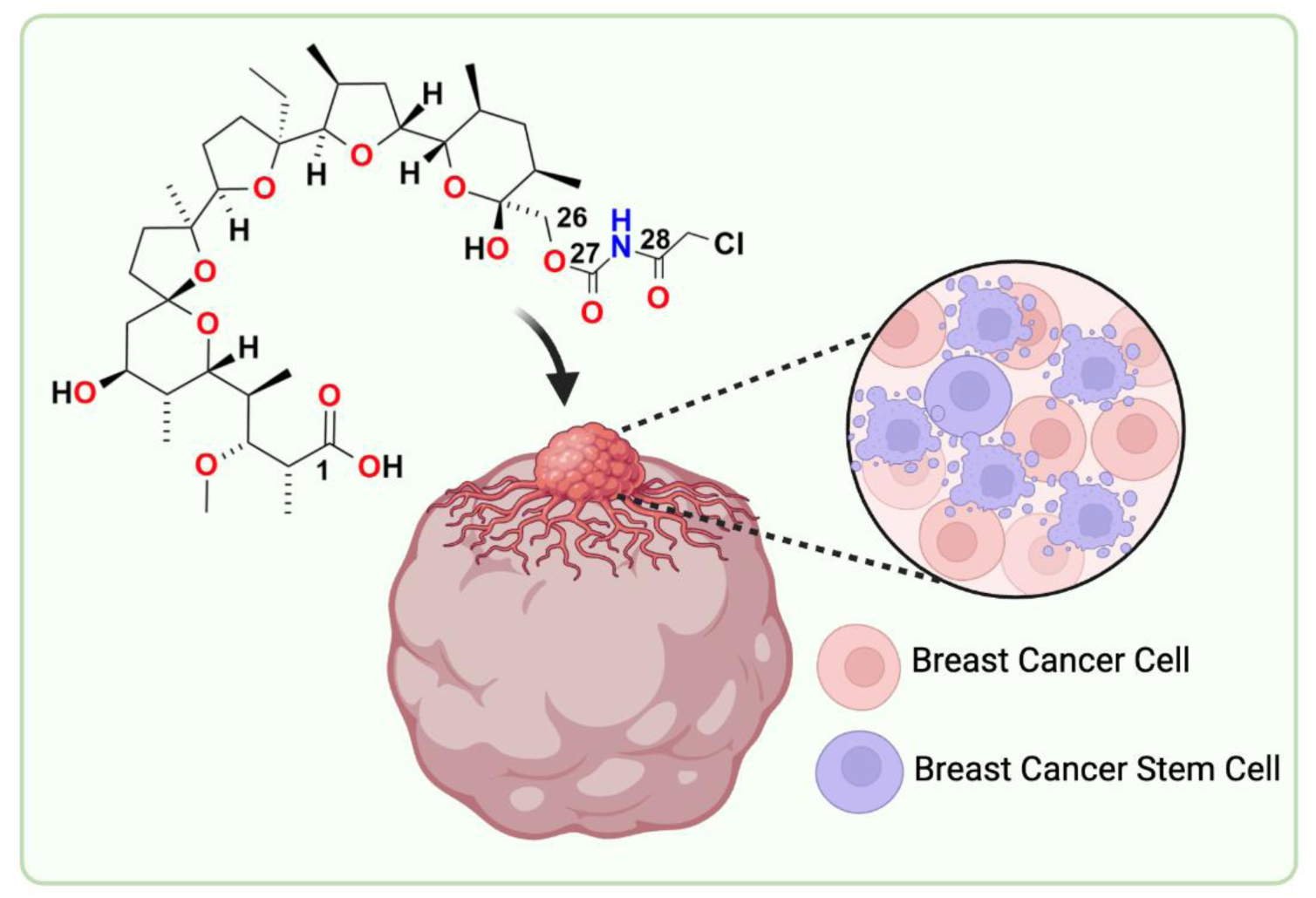

---

Breast cancer (BrCa) remains the most frequently diagnosed malignancy among women ^1^. Metastasis, characterized by the spread of cancer cells from the primary tumor to secondary sites, is the leading cause of BrCa-associated mortality, with a median survival of 8 to 36 months ^2^ ^3^. Primary metastatic sites for BrCa include lymph nodes, bones, lungs, liver, and brain ^2^ ^4^. A significant challenge in treating metastatic BrCa is the ability of cancer cells to remain dormant, rendering them resistant to anti-proliferative therapies ^2^ ^5^ ^6^. Consequently, there is an urgent need for therapeutic strategies to both prevent metastasis and to effectively treat metastatic tumors.

BrCa is clinically categorized into three subtypes based on receptor status: hormone-receptor (HR)-positive (ER+, PR+/-), HER2-positive (HER2+), and triple-negative BrCa (TNBC) (ER-, PR-, and HER2-) ^2^ ^7^ ^8^. Additionally, molecular profiling using the 50-gene PAM50 classifier revealed five intrinsic subgroups: luminal A, luminal B, HER2-enriched, basal-like and normal-like, underscoring the substantial molecular heterogeneity within and across BrCa subtypes ^2^ ^9^. HR+ cancers are often luminal A or luminal B. While luminal A tumors are typically HER2-negative, ER+, PR high, and Ki67 (a marker of cell proliferation) low, resulting in low-grade, slow-proliferating neoplasms, luminal B tumors are more aggressive, exhibiting variable PR and Ki67 levels and may be HER2-positive or HER2-negative ^2^ ^10^. TNBC, characterized by its aggressive nature, early metastasis, and poor prognosis, remains a particularly challenging subtype to treat ^11^.

Breast cancer stem cells (BCSCs) play a pivotal role in tumor initiation, progression, and metastasis ^12^. These cells drive long-term tumor growth and are responsible for recurrence and metastasis. BCSCs are most commonly identified by the expression of characteristic cell-surface markers, which vary across different subtypes of BrCa ^13^. In BrCa, high expression of CD44 with concomitant low expression of CD24 (CD44^+^/CD24^-/low^) is associated with cell proliferation and tumorigenesis ^13^ ^14^. Another functional marker of BCSCs, aldehyde dehydrogenase 1 (ALDH1), regulates CSC through two primary mechanisms. First, ALDH1 detoxifies harmful aldehydes (R-CHO) by converting them into their corresponding acids (R-COOH), protecting DNA, mitochondria, and other cellular components ^15^. This protective role is crucial for the self-renewal, survival, and proliferation of CSCs ^15^. ALDH1 may also shield CSCs from aldehyde-containing drug metabolites ^15^. Second, ALDH1 acts as a detoxifying enzyme that oxidizes retinol to retinoic acid, a molecule essential for early stem cell differentiation and widely used to characterize stemness ^13^ ^16^. Additionally, ALDH1 facilitates the conversion of retinal to retinoic acid, which binds to the retinoic acid receptor (RAR)/estrogen receptor α (ER α) complex, activating transcription of target genes such as c-Myc and cyclin D, thereby promoting CSC self-renewal, survival, and proliferation ^15^ ^17^. Notably, the polyether ionophore antibiotic salinomycin has shown the ability to selectively target BCSCs ^18^ ^19^. Another polyether ionophore, monensin (**MON**), produced by *Streptomyces cinnamonensis* has unique structural and functional properties that make it an intriguing candidate for anti-cancer therapy ^20^. **MON**’s cyclic structure, with protruding alkyl groups, renders it highly lipid-soluble, enabling it to traverse lipid bilayers and transport ions through passive diffusion ^20^ ^21^. **MON** exhibits a strong preference for Na^+^ over K^+^, making it particularly effective as an ion transporter ^22^.

**MON** is approved by the U.S. Food and Drug Administration (FDA) for veterinary applications as an anti-coccidiosis agent and has demonstrated a favorable safety profile through over 40 years of use in cattle and poultry feed ^23^ ^24^. Its widespread use underscores its biosafety and makes it as a strong candidate for drug repositioning. In recent years, **MON** has garnered attention as a potential anti-cancer agent, with demonstrated activity against renal cell carcinoma, lymphoma, myeloma, colon cancer, head and neck squamous cell carcinoma, glioblastoma, and prostate cancer and others ^25^ ^26^ ^27^ ^28^ ^29^ ^30^ ^31^. Remarkably, Ketola et al. reported that **MON** exhibited over 20-fold greater cytotoxic effect on malignant cell lines compared to non-malignant ones ^32^. **MON** has also shown potent activity against BrCa cells. Specifically, Gu et al. found that MON inhibited proliferation, migration, and the expression of MMP-2 and MMP-9 ^33^. It promoted apoptosis, as evidenced by increases in Bax, caspase-3, caspase-7, and caspase-9, a decrease in Bcl-2, and inhibition of UBA2 in luminal A, MCF-7 cells ^33^. Furthermore, **MON** liposomes have been shown to potentiate the cytotoxic activity of anticancer drugs in MCF-7/dox cells, overcoming drug resistance to doxorubicin, etoposide, and paclitaxel ^34^. Importantly, recent studies on TNBC cell lines, MDA-MB-231 and 4T1, have revealed that, similar to salinomycin, **MON** targets BCSCs, further supporting its therapeutic potential in BrCa ^35^.

In this study, we investigated whether compounds from a library of **MON** analogs exhibit enhanced anti-BCSC activity and improved selectivity for cancerous versus non-cancerous cells. To overcome the limitations of traditional 2D cell culture models, such as the lack of structural architecture and phenotypic drift during prolonged culturing, we employed a 3D cancer organoid model ^36^. This model better replicates the physiological and morphological characteristics of solid tumors like BrCa, offering a more reliable platform to evaluate responses to chemotherapy agents ^37^.

## 1. RESULTS AND DISCUSSION

### 1.1. Analogs design and synthesis

One of the key determinants of MON’s biological activity is its ability to complex metal ions. Therefore, through strategic chemical modifications—altering lipophilicity, the number of heteroatoms, and the balance of hydrogen bond donors and acceptors—we aimed to investigate how these structural changes influence its ion-complexing capacity and selectivity. Additionally, we explored whether such modifications could reduce cytotoxicity while preserving or even enhancing its efficacy. Incorporation of chemically labile moieties, such as esters, can be seen as a prodrug approach that masks **MON**’s ionophoric activity until it reaches its target site, potentially minimizing off-target effects and toxicity. Furthermore, chemical modification of **MON** could facilitate the discovery of a structurally diverse library of bioactive derivatives, offering new avenues to overcome resistance mechanisms not only in cancer therapy but also in antimicrobial applications. Over the years, diverse series of **MON** analogs have been synthesized and extensively studied with their biological evaluation reported in the literature. These include single modifications, such as esters at the C26 position (1–6), urethanes at the C26 (7–14), a lactone-derived cyclic analog (15), esters at the C1 position (16–21), tertiary amides at the C1 (22–25), and a cyclic carbonate at the C25 and C26 (26). Additionally, double-modified analogs incorporating a cyclic carbonate at the C25 and C26 in combination with either ester moieties at the C1 (27–33) or amide moieties at the C1 (34–37) have been also developed.

### 1.2. Activity of monensin and its analogs in mini-ring cell viability assay

The human TNBC basal-like mesenchymal cell line MDA-MB-231 is characterized by high expression of CD44^+^/CD24^-/low^ and elevated ALDH1 levels ^13^, both of which are characteristic markers of BCSCs. Consequently, this cell line was incorporated into all experiments in the present study. The anti-proliferative activity of **MON** and its synthetic analogs (**Fig. 1**) was evaluated using the MTT cell viability assay ^54^, adapted for the 3D mini-ring screen as previously described ^43^ ^44^ ^45^. In this model, cells are suspended in Matrigel and seeded around the rim of each well in a mini-ring format (**Fig. 2**), which enables more reliable cell-cell interaction and better recapitulates the tumor microenvironment than conventional cell monolayers, thereby providing a more reliable screening technique for drug discovery ^44^.

**Figure 1.**
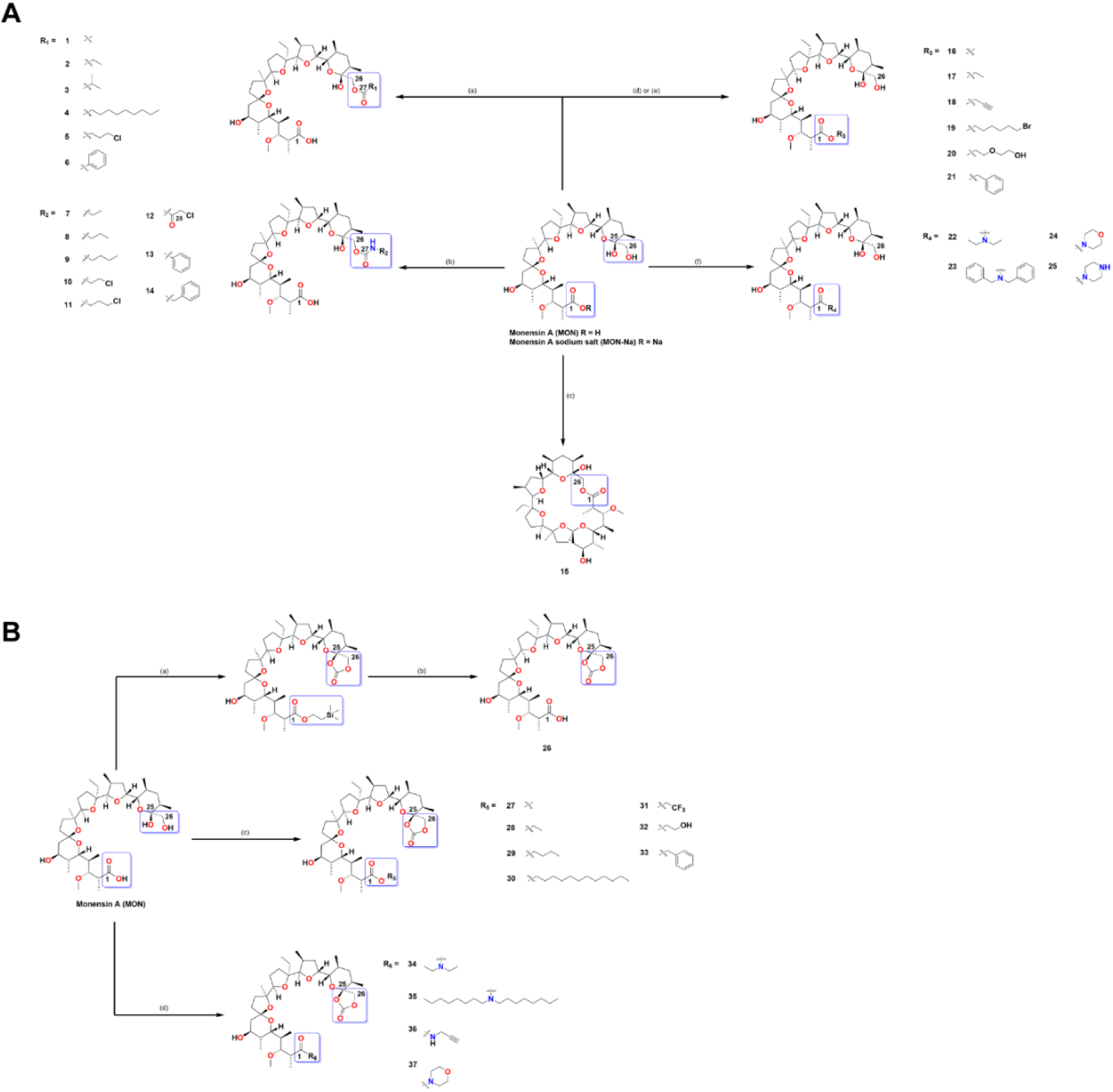
Synthesis of MON analogs. **A**. Reagents and conditions: (a) MON-Na (1 eq), DMAP (catalytic amount), respective acyl chloride (3 eq), Py, rt, 48 h; (b) MON (1 eq), respective isocyanate (0.95 eq), anh. toluene, rt, 14 days; (c) MON (1 eq), 2,2′-dipyridyl disulfide (2.5 eq), PPh_3_ (2.5 eq), toluene, reflux, 17 h; (d) for compounds 16, 17 and 20: MON (1 eq), DCC (1.5 eq), 4-pyrrolidinylpyridine (0.5 eq), *p*-TSA (0.25 eq), respective alcohol (10 eq), DCM, 0°C–2 h, then rt–22 h, (e) for compounds 18, 19 and 21: MON (1 eq), DBU (1.3 eq), respective akyl halide (3 eq), toluene, 80°C, 24 h; (f) MON (1 eq), DCC (1.2 eq), HOBt (0.5 eq), respective secondary amine (2.5 eq), DMF, 0 °C—1 h, then rt—24 h. **B**. Reagents and conditions: (a) MON (1 eq), TEA (5 eq), BTC (1.05 eq), 2-(trimethylsilyl)ethanol (20 eq), DMC, 0°C → rt; (b) 1. 1M TBAF in THF (3 eq), rt, 30 – 60 min;2. Na_2_CO_3_ aq. sol. wash; 3. H_2_SO_4_ aq. sol. (pH ≈ 1) wash; (c) MON (1 eq), TEA (5 eq), BTC (1.05 eq), respective alcohol (20 eq), DCM, 0 °C → rt; (d) MON (1 eq), TEA (5 eq), BTC (1.05 eq), respective amine (20 eq), DCM, 0° C → rt. Reaction time was determined by TLC.

**Figure 2.**
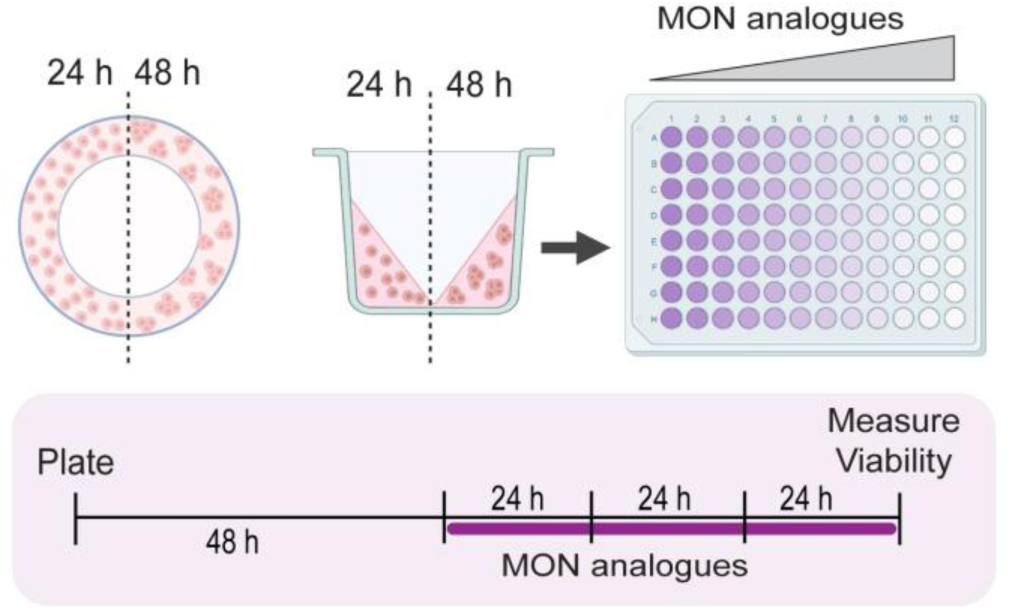
Schematic representation of compound response assessment using the 3D mini-ring cell viability assay. Cells seeded in the mini-rings were incubated for 48 h to allow growth and establishment. Following this, the cells were treated with compounds or control, and the medium containing the treatment was refreshed every 24 hours for a total of 72 hours. After the treatment period, cells were released from the mini-rings using dispase and subjected to the MTT assay, as described in the METHODS section.

The results, shown as IC_50_ ± SD, are presented in **Table 1**. The IC_50_ of parent **MON** against MDA-MB-231 cells was determined to be 650 nM, which is consistent with our previously observed IC_50_ value of 613 nM for this compound in a mini-ring cell viability assay against the U-118 MG glioblastoma cell line ^31^.

**Table 1.** Antiproliferative activity (IC_50_ ± SD) values of monensin (MON), and its analogs.

| Compound | MDA-MB-231 | MCF10A | SI |
| --- | --- | --- | --- |
| | $IC_{50}$ (nM) | | |
| MON | $650 \pm 459$ & | $221 \pm 102$ # | 0.3 |
| 1 | $1536 \pm 884$ | $441 \pm 177$ | 0.3 |
| 2 | $146 \pm 7.39^{***}$ | $61.5 \pm 17.0$ | 0.4 |
| 3 | $170 \pm 79.1^{***}$ | $67 \pm 44.1$ | 0.4 |
| 4 | $307 \pm 139$ | $26.1 \pm 17.0$ | 0.1 |
| 5 | $84.9 \pm 25.3^{****}$ | $1145 \pm 541$ | 13.5 |
| 6 | $304 \pm 113^*$ | $621 \pm 325$ | 2.0 |
| 7 | $214 \pm 151^{**}$ | $274 \pm 22.4$ | 1.3 |
| 8 | 262 ± 122** | 149 ± 55.0 | 0.6 |
| 9 | 236 ± 6.08** | 65.4 ± 19.0 | 0.3 |
| 10 | 232 ± 45.7** | 169 ± 80 | 0.7 |
| 11 | 813 ± 209 | 141 ± 68.6 | 0.2 |
| 12 | 122 ± 8.27 *** | 464 ± 160 | 3.8 |
| 13 | 370 ± 246 | 61.5 ± 31.6 | 0.2 |
| 14 | 1405 ± 664 | 45.2 ± 17.5 | 0.03 |
| 15 | 2884 ± 390 *** | 968 ± 289 | 0.3 |
| 16 | 9489 ± 1482 ** | N/A | N/A |
| 17 | 3904 ± 512 *** | N/A | N/A |
| 18 | 1070 ± 399 | N/A | N/A |
| 19 | >10000 | N/A | N/A |
| 20 | >10000 | N/A | N/A |
| 21 | 3546 ± 731 ** | 1035 ± 397 | 0.3 |
| 22 | >10000 | N/A | N/A |
| 23 | 2586 ± 504 ** | 2014 ± 202 | 0.8 |
| 24 | >10000 | N/A | N/A |
| 25 | >10000 | N/A | N/A |
| 26 | >10000 | N/A | N/A |
| 27 | 7233 ± 3241 * | N/A | N/A |
| 28 | 7708 ± 803 *** | N/A | N/A |
| 29 | >10000 | N/A | N/A |
| 30 | >10000 | N/A | N/A |
| 31 | >10000 | N/A | N/A |
| 32 | 2897 ± 1121 * | 1680 ± 1293 | 0.6 |
| 33 | >10000 | N/A | N/A |
| 34 | >10000 | N/A | N/A |
| 35 | >10000 | N/A | N/A |
| 36 | >10000 | N/A | N/A |
| 37 | >10000 | N/A | N/A |

IC<sub>50</sub> value is defined as the concentration of the compound which induces 50% growth inhibition.
SI (Selectivity Index) represents the ratio of a compound's cytotoxicity in non-cancerous cells to its efficacy in cancer cells, indicating its therapeutic window and specificity.
Values are given as mean ± SD (n = 4).
p values were calculated by comparing each treatment to MON using Welch's unpaired t-test. \*p ≤ 0.05, \*\*p ≤ 0.01, \*\*\* p ≤ 0.001, \*\*\*\*p ≤ 0.0001.
& The value is an average from five independent experiments (n = 18).

Esters and urethanes on the C26 atom exhibited the highest anti-BrCa cell activity (as determined by the lowest IC_50_ values) from the entire tested library of compounds. Esters R_1_ **2**, **3**, **5**, and **6** (**Fig. 1**) had significantly lower IC_50_ values of 146, 170, 85, and 304 nM, respectively, compared to the parent **MON** (650 nM). The IC_50_ values of urethanes R_2_ **7**, **8**, **9**, **10**, and **12** (**Fig. 1**) were 214, 262, 236, 232, and 122 nM, respectively, significantly lower than that of **MON**. Esters on the C1 atom (compounds R_3_ **16**-**21**, **Fig. 1**) were characterized by remarkably higher IC_50_ values in the mid-high micromolar range, with the lowest values of 3.9 and 3.5 µM recorded for analogs **17** and **21**, respectively. Among tertiary amides on C1 (compounds R_4_ **22**-**25**, **Fig. 1**), only analog **23** showed considerable activity with an IC_50_ of 2.6 µM. No other analog from this group showed remarkable activity at the tested concentrations. Among ester-carbonates (compounds **26** & R_5_ **27-33**, **Fig. 1**), compound **32** was characterized by the lowest IC_50_ of 2.9 µM, while two other compounds, **27** and **28**, had IC_50_ values in the range of 7 µM. Amide carbonates (compounds R_6_ **34**-**37**, **Fig. 1**) did not show the ability to kill at least 50% of cancer cells even at the highest concentration used (10 µM) (**Table 1**).

The Selectivity Index (SI) defines the ability of a compound to preferentially target tumor cells over non-cancerous cells. For this reason, the library of **MON** analogs was also tested against non-cancerous epithelial BrCa, MCF10A (**Table 1**). The SI of 0.3 for **MON** suggests that this compound may show preferential cytotoxicity against non-cancerous versus BrCa stem-like cells. Fortunately, some novel analogs, including compounds **5**, **6**, **7**, and **12**, had a favorable SI of >1, indicative of their potential higher activity toward cancer versus non-cancerous cells.

Analogs **5**, **6**, **12**, **13**, **15**, **21**, **23**, and **32**, as well as the parent **MON**, were selected for further studies. This selection was informed by low IC_50_ values coupled with high SI (for compounds **5**, **6**, and **12**), while also ensuring structural diversity across the chosen compounds.

### 1.3. Cell cycle analysis on BrCa organoids

To elucidate the mechanism by which **MON** and its analogs induce cell death in BrCa organoids, their impact on the cell cycle profile was assessed using flow cytometry. To better model the 3-dimensional (3D) structure of BrCa and address the limitations associated with cell monolayers, BrCa organoids were generated as described in the METHODS section and shown in **Fig. 3A**. These organoids were treated with 0.1% DMSO (control) or compounds **5**, **6**, **12**, **13**, **15**, **21**, **23**, and **32** at 5 x IC_50_ concentrations (**Table 1**) for 24 and 72 hours, with these time points chosen to assess early and late effects, respectively. After incubation, the organoids were dispersed into single cells, fixed, and stained with propidium iodide to assess DNA content and cell cycle phase. Cells with sub-G1 (<2N) DNA content were quantified as dead. The full set of representative cytograms is shown in **Fig. 3B**. A summary of MDA-MB-231 cells in different phases of the cell cycle, averaged over three independent biological replicates, is presented in **Fig. 3C**. Statistically significant increases in sub-G1 DNA content (control vs. compound treatment) were observed after 72 hours of incubation with all the compounds studied. **MON** and analogs **5**, **6**, **12**, **13**, **21**, and **23** induced DNA fragmentation as early as 24 hours. Notably, since the concentrations were adjusted to the respective 5 x IC_50_ values (**Table 1**), and analogs **5**, **6**, **12**, and **13** had significantly lower IC_50_ values than parent **MON**, these synthetic analogs were able to induce comparable DNA fragmentation at doses two to eight times lower than that of unmodified **MON**. The findings from this experiment, combined with the improved IC_50_ values and SIs, led us to select esters **5**, and **6**, and urethane **12** for further studies, with compound **13** excluded because of poor selectivity (**Table 1**).

**Figure 3.**
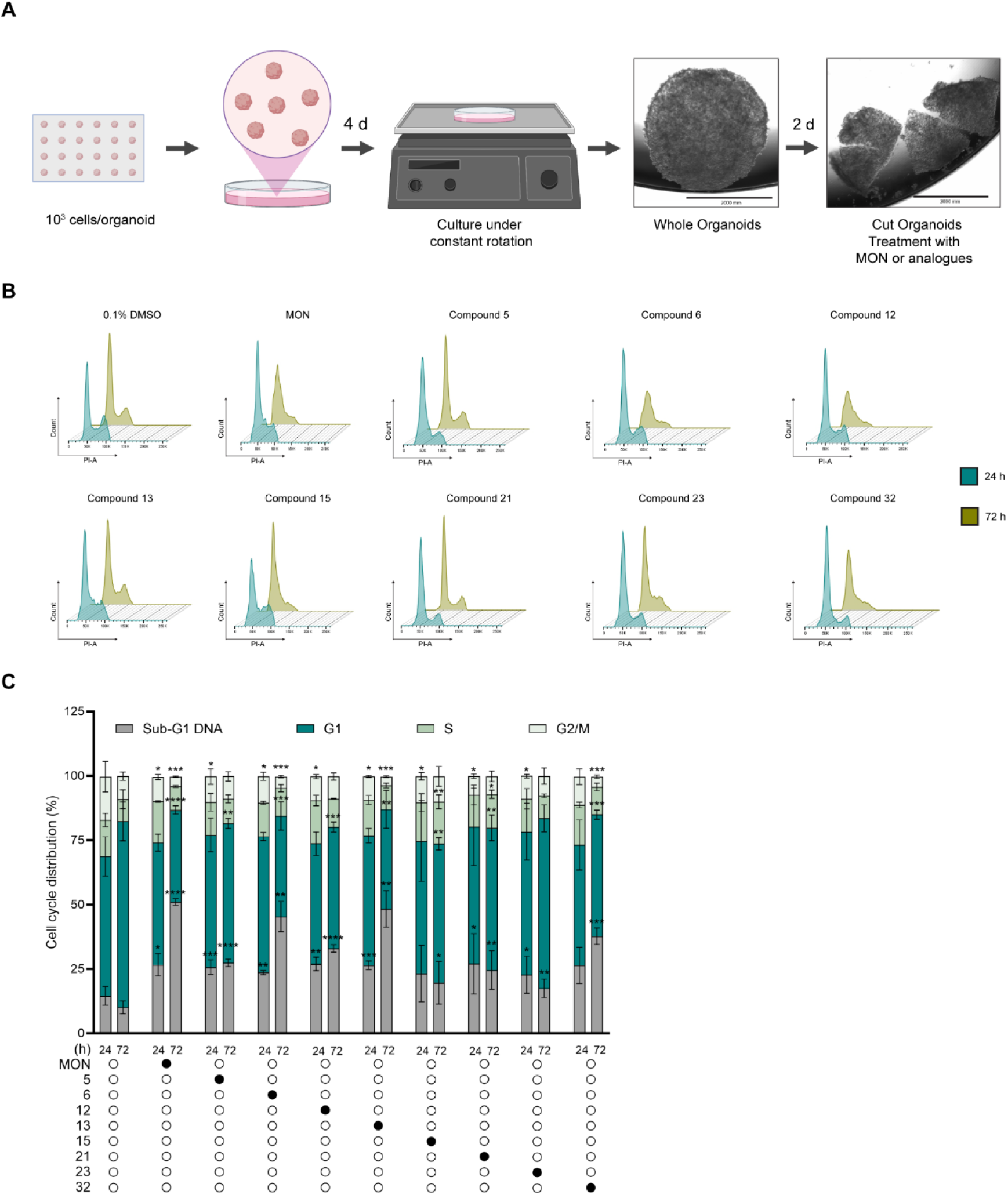
MON and its analogs induce DNA fragmentation in MDA-MB-231 breast cancer organoids. MDA-MB-231 organoids were treated with 0.1% DMSO (vehicle control), **MON** or analogs **5**, **6**, **12**, **13**, **15**, **21**, **23**, or **33** at concentrations equal to 5 x IC_50_ values (Table 1) for 24 or 72 hours. Following treatment, the organoids were subjected to propidium iodide staining and analyzed by flow cytometry. **A**. Schematic representation of MDA-MB-231 organoid generation and treatment. After two days of culturing under constant rotation, the organoids were cut to prevent the formation of a necrotic core and to maintain cellular proliferation. All subsequent experiments were conducted on these fragmented organoids. **B**. Representative cytograms. **C**. Distribution of cells across different phases of cell cycle and those with sub-G1 DNA content. Data are presented as mean ± SD (n = 3-6). Statistical significance is indicated relative to the DMSO control at the respective time points: each treatment at 24 h was compared to the DMSO control at 24 h, and each treatment at 72 h was compared to the DMSO control at 72 h: *p ≤ 0.05, **p ≤ 0.01, *** p ≤ 0.001, ****p ≤ 0.0001.

### 1.4. Monensin and compounds 6 and 12 attenuate migration of MDA-MB-231 cells

To evaluate whether **MON** or its analogs can inhibit the migratory properties of MDA-MB-231 cells, we performed a wound healing assay. Wounds were made on a confluent cell monolayer, and cells were allowed to migrate for up to 48 hours in the presence of 0.1% DMSO (control), **MON**, or compounds **5**, **6**, or **12** at concentrations equivalent to half of their respective IC_50_ values (**Table 1**), in order to affect migration without inducing cytotoxicity. Compound **6** significantly reduced the migration of MDA-MB-231 cells as early as 3 hours (93% vs. 87% of wound surface area for treatment vs. control, respectively) (**Fig. 4**). After 27h of incubation **MON**, along with compounds **6** and **12**, demonstrated significant inhibitory activity on the migration of MDA-MB-231 cells (**Fig. 4**). After 48 hours, analogs **6** and **12** significantly inhibited cell migration, with compound **6** showing the greatest effect (58% vs. 33% of wound surface area for treatment vs. control, respectively) (**Fig. 4**). Importantly, compound **6** and compound **12** were used in this assay at half and one fifth of the concentration of the unmodified **MON** respectively. Compound **5** did not exhibit significant anti-migratory properties in BrCa cells.

**Figure 4.**
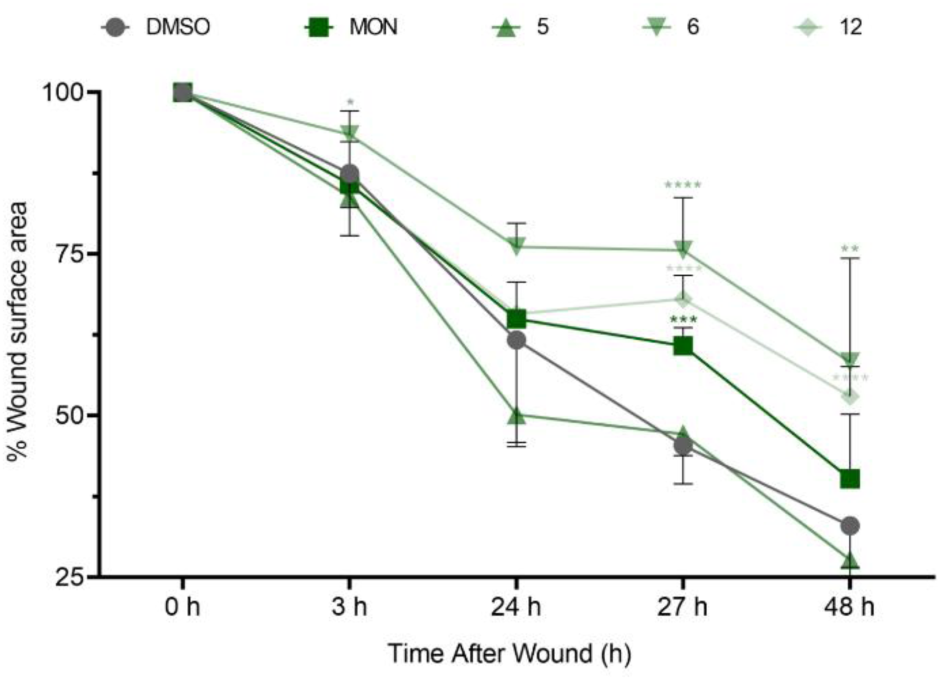
MON and analogs **6** and **12** attenuate MDA-MB-231 cell migration. A wound healing assay was performed on MDA-MB-231 cells treated with 0.1% DMSO (vehicle control), 325 nM **MON**, 42.5 nM compound **5**, 152.5 nM compound **6**, or 65 nM compound **12** for 48 hours. Wounds were created in a confluent cell monolayer, and images were captured at 0, 3, 24, 27, and 48 hours post-wounding. Three independent biological replicates were conducted. Data are presented as mean ± SD (n = 3). Statistical significance was determined by comparing each treatment group to the DMSO control (*p ≤ 0.05, **p ≤ 0.01, *** p ≤ 0.001, ****p ≤ 0.0001).

### 1.5. Treatment with monensin and compounds 6 and 12 downregulates the expression of SOX2

The activity of the transcription factor - sex-determining region Y (SRY)-box 2 (SOX2) has been linked to the maintenance of the undifferentiated state of CSCs in various tissues, including the breast ^55^. Increased levels of SOX2 have been reported in TNBC and are associated with poorer prognosis ^56^. Therefore, SOX2 silencing may be considered a novel therapeutic approach for TNBC ^56^. Pádua et al. previously demonstrated that **MON** exhibited selective toxicity toward gastric SORE6+ cells enriched with SOX2 ^57^, while Fang et al. reported significant downregulation of SOX2 in MDA-MB-231 and 4T1 BrCa cell lines following **MON** treatment ^35^.

In the present study (**Fig. 5A**), we show that, consistent with previous findings ^35^, **MON** significantly downregulated the expression of the SOX2 gene in MDA-MB-231 BrCa cells compared to the untreated control. Similarly large decreases were observed for **MON** analogs, compounds **6** (an 80% decrease) and **12** (an 89% decrease) (**Fig. 5A**). In contrast, treatment with compound **5** led to only a 20% decrease in SOX2 expression that was not statistically significant (**Fig. 5A**).

**Figure 5.**
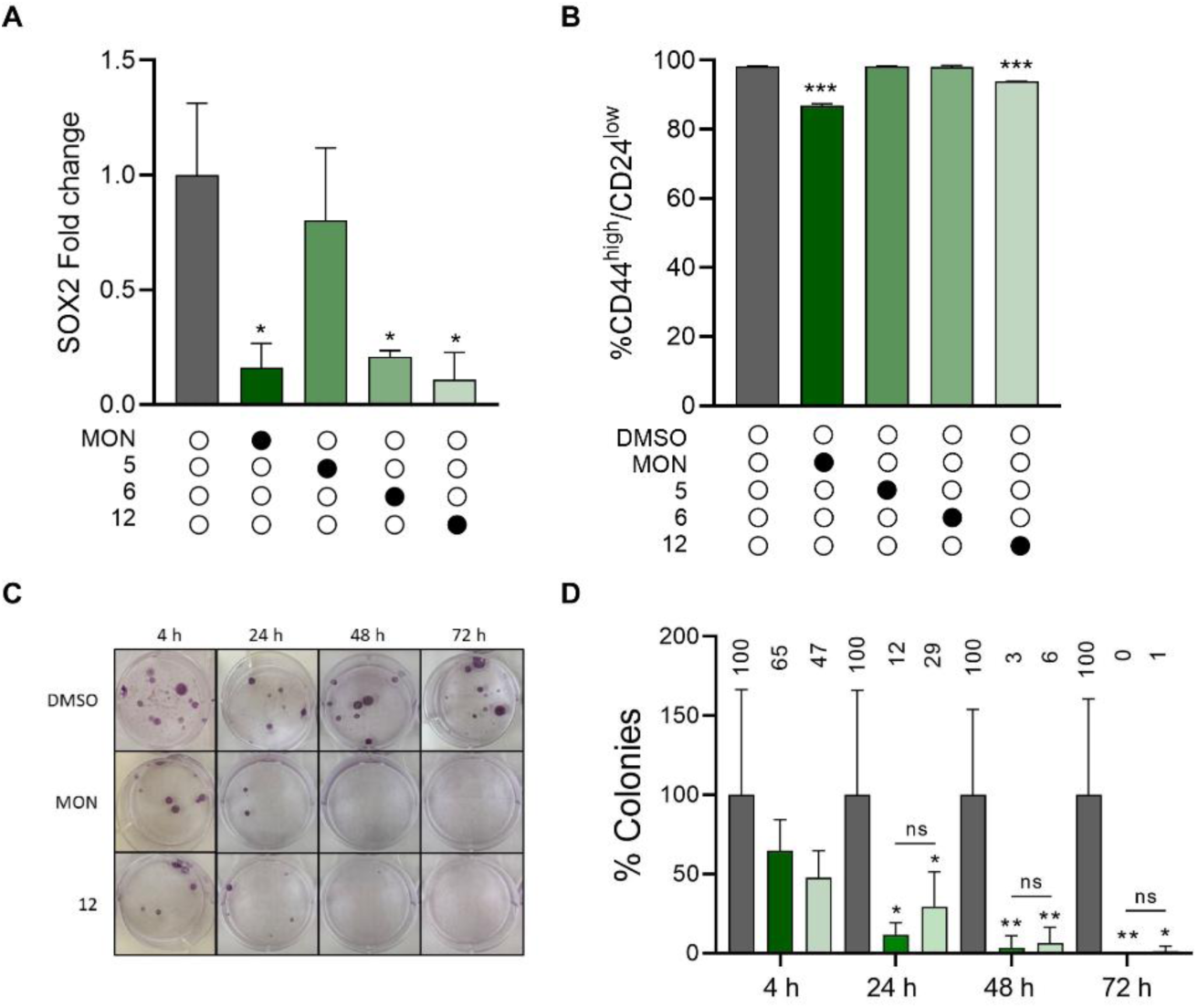
MON and analogs show anti-breast cancer stem-like cell activity in the MDA-MB-231 cell line. A. The effect of a 72-hour treatment with MON (325 nM), or analog 5 (45.5 nM), 6 (152.5 nM), or 12 (65 nM) on SOX2 mRNA levels was assessed. Bar graphs represent the mean ± SD of three biological replicates (n = 3). B. MON and analog 12 downregulate the expression of CD44^high^/CD24^low^ stem-like populations in MDA-MB-231 cell line. MDA-MB-231 cells were triple-stained with anti-CD44-PE, anti-CD24-FITC, and propidium iodide (PI). The ratio of viable PI-negative CD44^high^/CD24^low^ MDA-MB-231 cells following treatment with 0.1% DMSO (vehicle control), 325 nM MON, 42.5 nM compound 5, 152.5 nM compound 6, or 65 nM compound 12 for 72 hours. Data are presented as mean ± SD (n = 3). C. Effect of MON and analog 12 on the colony formation potential of MDA-MB-231 cells. MDA-MB-231 cells were treated with 0.1% DMSO (vehicle control), 650 nM MON, or 130 nM compound 12 for 4, 24, 48 and 72 hours. Following treatment, cells were harvested and re-seeded into 6-well plates at a density of 250 cells per well. After 21 days, colonies were fixed and stained with a 0.5% crystal violet solution in methanol. Colonies were manually counted for each well and normalized to the control. D. Quantification of colony numbers. Data are presented as mean ± SD (n = 6). Numerical percentage values are displayed above each bar. Statistical significance was determined by comparing each treatment group to the DMSO control: *p ≤ 0.05, **p ≤ 0.01, *** p ≤ 0.001.

### 1.6. Monensin and compound 12 reduce the CD44^+^/CD24^-/low^ stem-like cell population in MDA-MB-231 cells

We next examined the effect of **MON** and compounds **5**, **6**, and **12** on the CD44^+^/CD24^-/low^ stem-like population within MDA-MB-231 cells. Quantification of viable CD44^+^/CD24^-/low^ populations relative to the DMSO control is shown in **Fig. 5B**. As previously reported ^35^, **MON** induced a modest (∼11%), but statistically significant (p < 0.001) reduction in the percentage of viable CD44^+^/CD24^-/low^ cells, accompanied by an increase in the CD44^+^/CD24^+^ population (**Fig. 5B**). Similarly, compound **12** induced a small (∼6%), but statistically significant (p < 0.001) decrease in the percentage of viable CD44^+^/CD24^-/low^ cells (**Fig. 5B**). In contrast, analogs **5** and **6** showed no perceptible effect on the percentage of viable CD44^+^/CD24^-/low^ cells (decreases of <1%; p > 0.05) (**Fig. 5B**). Notably, since the concentrations were adjusted to half of the IC_50_ values (**Table 1**), compound **12** was able to produce about half the effect as the parent MON, but at five times lower dose.

### 1.7. Monensin and compound 12 reduce colony formation potential of MDA-MB-231 cell line

To determine the impact of **MON** and compound **12** on the self-renewal of MDA-MB-231 cells ^58^, we performed a clonogenic assay. This *in vitro* assay is used to evaluate whether an anti-cancer agent can inhibit the survival of tumor-initiating cells following treatment ^59^. It measures the capacity of single cells to divide and form colonies after a short exposure to a cytotoxic agent ^59^ ^60^. Briefly, MDA-MB-231 cells were plated in 6-well plates and treated for 4, 24, 48, or 72 hours with concentrations equal to IC_50_ values (see **Table 1**) of **MON**, compound **12**, or 0.1% DMSO (vehicle control). After treatment, cells were re-plated at low densities (250 cells per well in 6-well plates) and allowed to grow for 21 days. The results are shown in **Fig. 5C-D**. **MON** and compound **12** significantly reduced the colony-forming ability of MDA-MB-231 cells as early as 24 hours post-treatment, with further reductions observed at 48 and 72 hours (**Fig. 5D**). Together, these data indicate that **MON** and compound **12** significantly inhibit colony formation and, thus, the self-renewal and tumor-initiating capacity of MDA-MB-231 BrCa cells.

### 1.8. Monensin and compound 12 upregulate TIMP2 in MDA-MB-231 cells

After confirming that **MON** and compound **12** show activity against BrCa stem-like cells, molecular mechanisms underlying this effect were investigated. To this end, we conducted both proteomic and phospho-proteomic analyses and identified differentially regulated protein and phospho-proteins (FDR-adjusted p-value < 0.05 and an absolute fold change > 2) after **MON** or compound **12** treatment in MDA-MB-231 cells (**Table S1**, Supplementary Material). Using the Reactome Pathway Database, pathways significantly modulated by treatment with **MON** or compound **12** in MDA-MB-231 cells were identified. The top significantly differentially expressed pathways for **MON** or compound **12** are summarized in **Fig. 6 A-H**. Several of the top ten upregulated pathways by **MON** overlapped with those upregulated by compound **12**, including the Unfolded Protein Response (UPR), IRE1 alpha activation of chaperones, regulation of insulin-like growth factor (IGF) transport and uptake by insulin-like growth factor binding proteins (IGFBPs), platelet degranulation, TP53 regulation of transcription of death receptors and ligands, post-translational protein phosphorylation, extracellular matrix (ECM) organization, serine biosynthesis, and interleukin-10 signaling (**Fig. 6 A and C**). These shared pathways suggest similarities in the mechanisms of action of the two compounds. Additionally, both **MON** and compound **12** significantly downregulated several pathways, including complex I biogenesis, the citric acid (TCA) cycle and respiratory electron transport, metabolism of proteins, and rRNA processing in mitochondria (**Fig. 6 B and D**), further supporting similar mechanisms of action.

**Figure 6.**
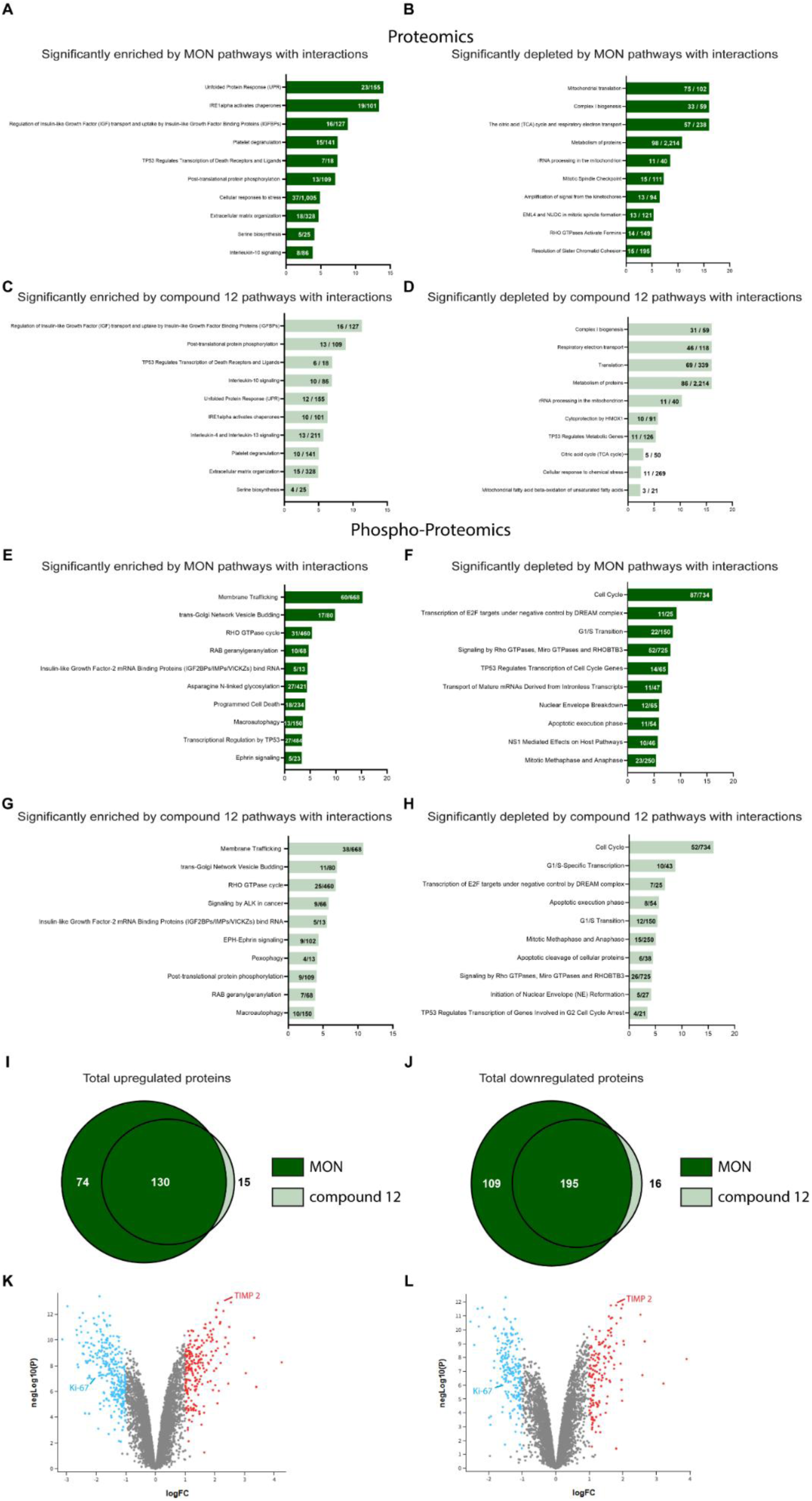
Proteomics and phosphoproteomics analyses identify key pathways upregulated and downregulated by **MON** and compound **12**, with TIMP2 significantly upregulated by both treatments. MDA-MB-231 cells were treated with 0.1% DMSO (vehicle control), 325 nM **MON**, or 65 nM compound **12** for 72 hours. **A**. Bar graph showing the 10 most significantly upregulated pathways in the proteomics analysis of **MON**-treated cells; **B**. Bar graph showing the 10 most significantly downregulated pathways in the proteomics analysis of **MON**-treated cells; **C**. Bar graph showing the 10 most significantly upregulated pathways in the proteomics analysis of compound **12**-treated cells; **D**. Bar graph showing the 10 most significantly downregulated pathways in the proteomics analysis of compound **12**-treated cells; **E**. Bar graph showing the 10 most significantly upregulated pathways in the phosphoproteomics analysis of **MON**-treated cells; **F**. Bar graph showing the 10 most significantly downregulated pathways in the phosphoproteomics analysis of **MON**-treated cells; **G**. Bar graph showing the 10 most significantly upregulated pathways in the phosphoproteomics analysis of compound **12**-treated cells; **H**. Bar graph showing the 10 most significantly downregulated pathways in the phosphoproteomics analysis of compound **12**-treated cells; **I**. **J**. Venn diagrams showing a total number of significantly upregulated (**I**), and downregulated (**J**) proteins by **MON** and compound **12**; **K**, **L**. Volcano plots highlighting proteins significantly downregulated (blue) or upregulated (red) in MDA-MB-231 cells treated with **MON** (**K**) or compound **12** (**L**) for 72 hours.

Phospho-proteomics analysis also revealed **MON** and compound **12** treatment induced overlapping enriched pathways, such as membrane trafficking, trans-Golgi network vesicle budding, RHO GTPase cycle, RAB geranylgeranylation, IGF2BPs/IMPs/VICKZ RNA binding, macroautophagy, and ephrin signaling (**Fig. 6 E and G**). The analysis of significantly downregulated pathways in the phospho-proteomics data further revealed shared effects of **MON** and compound **12** on cell cycle regulation, transcription of E2F targets under DREAM complex control, G1/S transition, signaling by Rho GTPases, Miro GTPases and RHOBTB3, TP53 regulation of cell cycle gene transcription, nuclear envelope breakdown and reformation, apoptotic execution phase, and mitotic metaphase and anaphase (**Fig. 6 F and H**). These findings suggest potential similarities in the signaling mechanisms by which **MON** and compound **12** target BCSCs.

Pathways associated with cell cycle progression were among the most affected by treatment (**Fig. 6**). The impact of **MON** treatment on cell cycle progression is well-documented across various cancer cell lines. For instance, **MON** has been reported to downregulate Cyclin D1 in SKOV3 ovarian cancer cells ^61^, PC-3 prostate cancer cells in a concentration-dependent manner, ^62^ and SNU-C1 colon cancer cells in a time-dependent manner ^63^. Our proteomics studies in MDA-MB-231 cells revealed similar findings, with **MON** (adj. p < 0.0001, Log fold change (FC) =-0.58) and analog **12** (adj. p < 0.001, LogFC =-0.43) downregulating Cyclin D1. Additionally, the same authors observed Cyclin E downregulation in PC-3 prostate cancer cells ^62^, which aligns with our results for **MON** (adj. p < 0.0001, LogFC =-0.88) and compound **12** (adj. p < 0.0001, LogFC =-0.77). **MON** has also been reported to induce concentration-or time-dependent downregulation of cyclin-dependent kinases (CDK) 2, 4, and 6 in PC-3 prostate ^62^ and SNU-C1 colon ^63^ cancer cells. Similarly, we observed a comparable effect in MDA-MB-231 cells, with **MON** significantly reducing CDK2 (adj. p < 0.0001, LogFC =-0.54), CDK4 (adj. p < 0.01, LogFC =-0.24), and CDK6 (adj. p < 0.01, LogFC =-0.18). Compound **12** also downregulated CDK2 (adj. p < 0.0001, LogFC =-0.45), CDK4 (adj. p < 0.05, LogFC =-0.18), and CDK6 (adj. p < 0.05, LogFC =-0.13).

**MON** has been previously reported to induce apoptosis in various cancer cell lines. In MCF-7 BrCa cells ^33^ Gu et al. demonstrated that **MON** upregulated caspase-7 and caspase-9 ^33^, which was consistent with our findings, where **MON** significantly increased caspase-7 fragments (adj. p < 0.001, Log FC =0.36) and caspase-9 fragments (adj. p < 0.001, logFC = 0.3). Compound **12** also upregulated caspase-9 (adj. p < 0.001, logFC = 0.32).

Our previous studies on glioblastoma organoids and other research on SKOV3 ovarian cancer cells indicated that **MON** significantly downregulates STAT3 ^45^ ^61^. This effect was also observed in our proteomics analysis of MDA-MB-231 for both **MON** (adj. p < 0.01, LogFC =-0.25) and compound **12** (adj. p < 0.05, LogFC =-0.18).

**MON** has been shown to downregulate multiple proteins involved in EMT. In SKOV3 cells, Yao et al. observed that **MON** induced concentration-dependent downregulation of vimentin, as shown by immunoblot analysis ^64^. Our proteomics data corroborate this, showing vimentin downregulation by **MON** (adj. p < 0.05, LogFC =-0.4), but not by analog **12**. Additionally, the same study reported MON-induced upregulation of claudin ^64^, which we confirmed in our dataset, with **MON** (adj. p < 0.01, LogFC = 0.6) and compound 12 (adj. p < 0.05, LogFC = 0.49) upregulating claudin-4.

Ochi et al. demonstrated that **MON** suppressed TGF-β induced vimentin upregulation in HCC827 and A549 cells and inhibited ZEB1 upregulation in H1975, A549 and PANC-1 cells ^65^. Our proteomics data similarly show significant downregulation of ZEB1 by **MON** (adj. p < 0.0001, LogFC =-0.4) and compound **12** (adj. p < 0.01, Log FC =-0.21).

Tumova et al. first reported that **MON** reduced β-catenin levels in colorectal carcinoma cells with deregulated Wnt signaling ^66^. Our proteomics analysis in MDA-MB-231 cells revealed similar effect, with **MON** (adj. p < 0.001, LogFC =-0.6) and compound **12** (adj. p < 0.01, LogFC =-0.4) reducing β-catenin levels. The same study found that **MON** promoted LRP6 degradation in Wnt3a-activated HEK293 cells ^66^. We observed comparable results in MDA-MB-231 cells, where **MON** (adj. p < 0.01, LogFC =-0.71) and compound **12** (adj. p < 0.001, LogFC =-0.92) downregulated LRP6.

In line with previous studies on CaSki and HeLa cervical cancer cells, we observed significant downregulation of phosphorylated β-catenin at Ser552 by **MON** (adj. p < 0.05, LogFC = 0.87), though this effect was not observed with compound **12** ^67^.

Wang et al. reported that **MON** (10 mg/kg) to downregulated proliferating cell nuclear antigen (PCNA) in a human pancreatic cancer xenograft model and reduced STAT1 levels in Panc-1 cells ^68^. Deng et al. also reported PCNA downregulation in an ovarian cancer xenograft model ^61^. Our proteomics data confirmed PCNA downregulation in MDA-MB-231 cells treated with **MON** (adj. p < 0.0001, LogFC =-0.37) or compound **12** (adj. p < 0.001, LogFC =-0.28). STAT1 was also downregulated by **MON** (adj. p = 0.055, LogFC =-0.14) and compound **12** (adj. p < 0.05, LogFC =-0.19). Furthermore RAF1, which was significantly repressed in Panc-1 and MiaPaCa2 cells in the same study, was similarly downregulated in our proteomics analysis following treatment with **MON** (adj. p < 0.0001, LogFC =-0.56) and compound **12** (adj. p < 0.001, LogFC =-0.46).

Overall, 130 proteins were significantly upregulated by both **MON** and compound **12** (**Fig. 6 I**), while 74 and 15 proteins were uniquely upregulated by **MON** or compound **12**, respectively. Similarly, 195 proteins were commonly downregulated by both treatments (**Fig. 6 J**), whereas 109 and 16 proteins were uniquely significantly downregulated by **MON** and compound **12**, respectively. These differentially expressed proteins may help explain the differences occasionally observed in the biological activities of these two molecules. Among the significantly downregulated proteins, Arylsulfatase B (ARSB) has been implicated in glioma progression by regulating M2 macrophage infiltration and the JAK2/STAT3 pathway ^69^. The downregulation of this protein by compound **12** (adj. p < 0.0001, LogFC =-1.12) may contribute to its superior biological activity compared to **MON** in our study, potentially explaining its enhanced therapeutic effect.

Of the identified pathways, extracellular matrix (ECM) organization was of particular interest (**Fig. 6 A and C**), as ECM remodeling is a critical event promoting cancer invasion and metastasis ^70^. ECM components such as integrins, collagen and fibronectin play key roles in cell adhesion, invasion and metastasis ^70^. Tissue inhibitor of metalloproteinase 2 (TIMP2), a major ECM constituent in normal tissues, was among the most overexpressed proteins in the ECM organization pathway following treatment, with a ∼6-fold increase after **MON** treatment and a 4-fold increase after compound **12** treatment (**Fig. 6 K and L**). TIMP2, an endogenous inhibitor of matrix metalloproteinase-2 (MMP-2), regulates tumor invasion by modulating MMP-2 activity ^71^ as well as anti-angiogenic and anti-tumor effects through MMP-independent mechanisms, including inhibition of growth signaling pathways ^71^ ^72^ ^73^ ^74^. Previous studies have demonstrated that TIMP2 suppresses TNBC growth and metastasis by modulating epithelial-to-mesenchymal transition, vascular normalization, and metastatic signaling pathways ^75^. These findings highlight TIMP2’s potential as both a direct and systemic anti-tumor and metastasis suppressor, with implications for clinical management of BrCa progression ^75^. Furthermore, TIMP2 overexpression has been shown to reduce invasion of endothelial and tumor cells *in vitro* and *in vivo* ^73^ ^76^ ^77^ ^78^ ^79^. Thus, compounds that upregulate TIMP2 may play a central role in anti-tumor and anti-invasive activity, positioning TIMP2 modulation as a potential molecular target for anti-metastatic therapies ^71^.

The upregulated expression of TIMP2 by **MON** and compound **12** identified through proteomics analysis was validated via immunoblotting (**Fig. 7 A-B**). To determine whether TIMP2 upregulation by **MON** and compound **12** is applicable across other BrCa subtypes and cancers, we examined TIMP2 expression in additional models. Both compounds significantly increased TIMP2 levels in MDA-MB-468 cells (a basal-like epithelial subtype of BrCa) (**Fig. 7 C-D**) and SK-MEL-5 melanoma cells (**Fig. 7 E-F**) which are also known to metastasize to the brain^80^.

**Figure 7.**
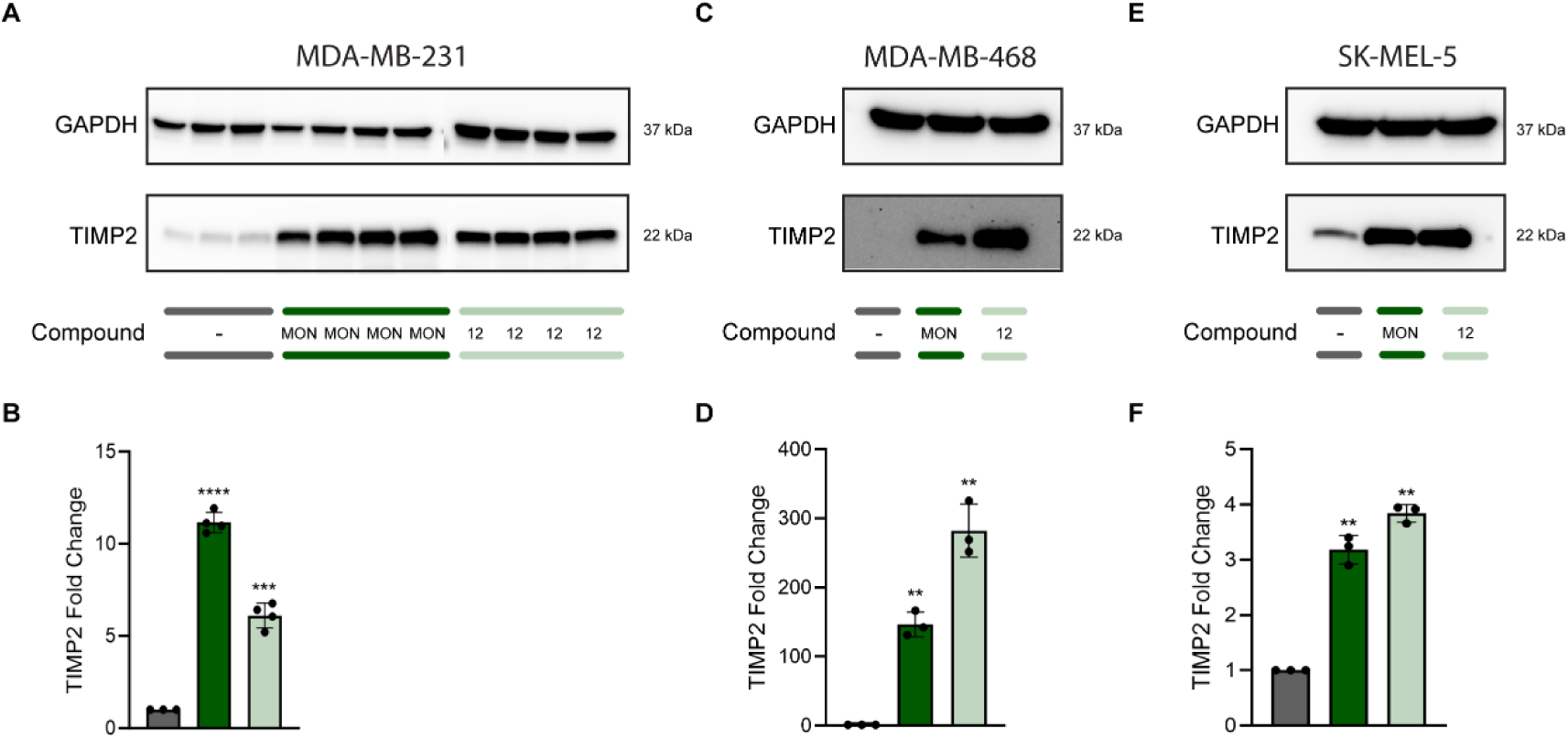
MON and compound 12 upregulate TIMP2 in MDA-MB-231, MDA-MB-468, and SK-MEL-5 cell lines. **A**. Immunoblot analysis of TIMP2 in MDA-MB-231 cell lysates, with GAPDH used as a loading control; **B**. Bar diagram showing fold changes in TIMP2 levels normalized to GAPDH in MDA-MB-231 cells; **C**. Immunoblot analysis of TIMP2 in MDA-MB-468 cells treated with 0.1% DMSO (vehicle control), 469 nM MON, or 332 nM compound 12; **D**. Bar diagram showing fold changes in TIMP2 levels normalized to GAPDH in MDA-MB-468 cells; **E**. Immunoblot analysis of TIMP2 in SK-MEL-5 cells treated with 0.1% DMSO (vehicle control), 250 nM MON, or 103 nM compound 12; **F**. Bar diagram showing fold changes in TIMP2 levels normalized to GAPDH in SK-MEL-5 cells. Images were quantified by measuring band intensity using ImageJ software. Data are presented as mean ± SD (n = 3–4). Statistical significance vs DMSO control: **p ≤ 0.01, ***p ≤ 0.001, ****p ≤ 0.0001.

### 1.9. MON and compound 12 show anti-metastatic BrCa activity in a 3D HOSTBO organoid model

Brain and leptomeningeal metastasis continue to be highly morbid complications of advanced BrCa, significantly impacting both quality of life and overall survival ^81^. To investigate the activity of **MON**, and its most potent analog, compound **12**, in metastatic BrCa cells within a human brain microenvironment, we developed hiPSCs-derived cerebral organoids (COs) as we previously described ^45^. These were co-cultured with MDA-MB-231 cells stably expressing red fluorescent protein (RFP) to generate human Hybrid Organoid System: Tumor-in-Brain Organoid (HOSTBO).

Briefly, the human COs were generated from hiPSCs constitutively expressing green fluorescent protein (GFP) under control of the ACTB promoter, using a four-stage protocol. This protocol involves CO formation through intermediate embryoid bodies, followed by the expansion of neuroepithelia ^45^. COs are 3D culture models characterized by structural organization and cellular composition resembling the developing human brain ^82^ ^83^. Following a maturation period, COs were sectioned and characterized using H&E staining, as well as immunofluorescence staining (**Fig. S1**). Routine H&E staining demonstrated embryonal cells with high nuclear-to-cytoplasmic ratios arranged in true rosettes as well as distributed within a neuropil-like background, consistent with developing brain (**Fig. S1A**). Embryonic central nervous system structures, such as a proliferative zone of neural stem cells (Nestin+) (**Fig. S1B**), neurons (TUJ1 +), a primitive ventricular system (SOX2 +/Ki67 +), an intermediate zone (TBR2 +), and a cortical plate (MAP2 +) were highlighted using immunofluorescence staining (**Fig. S1B**). Positive staining for the apical tight junction protein N-cadherin further indicated regulated development, suggesting a polarity like that found within the neural tube of the embryonic neural plate (**Fig. S1B**). These morphological and immunofluorescence findings confirm that the generated COs reflect the differentiation and development stage of an early-stage human fetal brain.

To model BrCa metastasis to the brain, individual, fully formed eGFP-expressing COs were co-cultured with 100,000 RFP-labeled MDA-MB-231 cells as previously optimized ^45^. After seven days of co-culture and substantial tumor formation, HOSTBOs were treated with 0.1% DMSO (control), **MON**, or compound **12**.

To assess the potency of **MON** and compound **12** on BrCa cells metastatic to the brain within a normal brain environment, HOSTBOs were treated as described in METHODS, sectioned to reveal RFP-labeled MDA-MB-231 cells internalized within the eGFP-labeled CO and analyzed for Ki-67, SOX2, cleaved caspase-3, and cleaved PARP expression (**Fig. 8 A-H**).

**Figure 8.**
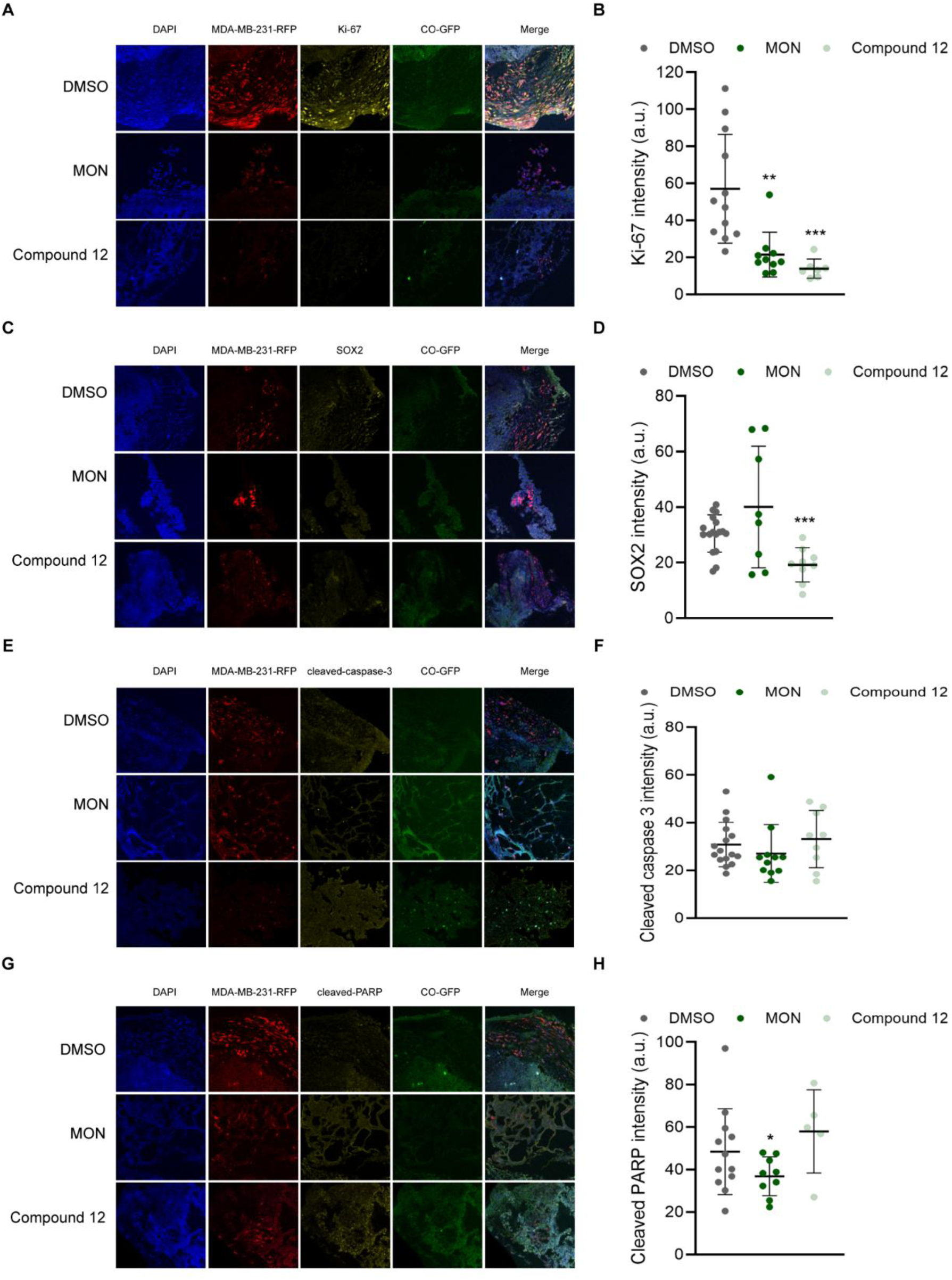
The response of HOSTBOs to treatment with **MON** or compound **12**. HOSTBOs were established by culturing eGFP-expressing COs with RFP-expressing MDA-MB-231 cells for 7 days and were subsequently treated with 0.1% DMSO (vehicle control), 1300 nM **MON** or 260 nM compound **12** for 4 days. Representative immunofluorescence images at 20X magnification are shown for (**A**) Ki-67, (**C**) SOX2, (**E**) cleaved caspase-3, and (**G**) cleaved-PARP. Quantification of staining intensity for (**B**) Ki-67, (**D**) SOX2, (**F**) cleaved caspase-3, and (**H**) cleaved-PARP. Data are presented as mean ± SD of technical replicates from two independent biological replicates. Statistical significance vs DMSO control: *p ≤ 0.05, **p ≤ 0.01, ***p ≤ 0.001.

Both **MON** and compound **12** significantly reduced the expression of the proliferation marker Ki-67 (**Fig. 8 A-B**), which is noteworthy, as inhibiting cancer cell proliferation is crucial for limiting tumor growth and progression. Notably, Ki-67 was also identified as one of the proteins significantly downregulated by both treatments in the proteomics analysis (**Fig. 6 K-L**). Additionally, compound **12** significantly downregulated SOX2 expression (**Fig. 8 C-D**), which is consistent with previous findings in cell monolayer (**Fig. 5A**). Although SOX2 was not detected in our proteomics data (**Fig. 6**), we observed a decreased expression of SOX9 following the treatment with **MON** (adj. p < 0.001, LogFC =-0.61) and compound **12** (adj. p < 0.01, LogFC =-0.45). SOX9 has been identified as a “master regulator” of BrCa cell survival and metastasis, as its loss has been shown to induce apoptosis ^84^. SOX9 directly binds to the promoters of genes involved in EMT including vimentin, claudin-1, β-catenin, and ZEB1 and its knockdown suppresses the expression of these genes ^84^.

Neither **MON** nor compound **12** induced cleavage of caspase-3 (**Fig. 8 E-F**), in contrast to the findings of Fang et al., who reported MON-induced caspase-3 cleavage in MDA-MB-231 and 4T1 BrCa cell lines ^35^. This discrepancy may be due to differences between cell monolayer and 3D organoid models, as well as the influence of the normal brain microenvironment. Further detailed mechanistic studies are needed to definitively determine whether these compounds induce metastatic BrCa cell death in a caspase-dependent or-independent manner. Similarly, no statistically significant effect on cleaved PARP expression was observed with compound **12** treatment (**Fig. 8 G-H**). In contrast, **MON** slightly but significantly downregulated cleaved PARP, differing from the observations of Fang et al. in MDA-MB-231 and 4T1 BrCa monolayers ^35^ and Kim et al. in PC-3 human prostate cancer cells ^85^. Once again, these differences may be attributed to variations between cell monolayers and 3D organoids, as well as the influence of the human brain microenvironment.

## 2. METHODS

### 2.1. Compounds

All compounds used in this study have been previously reported, along with their full analytical characterization. For reference, the original articles provide comprehensive experimental procedures, including NMR, MS, IR, and other relevant analytical data regarding compounds 1-6, 15-21 ^38^, 7-14 ^39^, 22-25 ^40^, and 26-37 ^41^. Readers are encouraged to consult these sources for further details.

### 2.2. Cells and culturing conditions

The following cell lines were used in this study: human breast adenocarcinoma MDA-MB-231 and MDA-MB-468, MDA-MB-231 stably expressing two separate reporters - firefly luciferase (Luc 3) and red fluorescent protein (RFP) (cat. no SC041, GenTarget Inc., San Diego, CA, USA), human breast epithelial MCF 10A, and human malignant melanoma SK-MEL-5. MDA-MB-231 and MDA-MB-468 cell lines were cultured in DMEM/Ham’s Nutrient Mixture F12 (1:1) (cat. no. 10-090-CV, Corning, Manassas, VA, USA) supplemented with 5% (v/v) heat-inactivated fetal bovine serum (FBS) (cat. no. FP-0500-A, Atlas Biologicals, Fort Collins, CO, USA). MCF 10A cell line was cultured in Mammary Epithelial Cell Basal Medium (cat. no. C-21215, Promo Cell, Heidelberg, Germany) supplemented with Mammary Epithelial Cell Supplement Pack (cat. no. C-39110, Promo Cell, Heidelberg, Germany), 100 µg/ml gentamicin (cat. no. G1397, Sigma-Aldrich, Deisenhofen, Germany), and 0.05 µg/ml amphotericin B (cat. no. A2942, Sigma-Aldrich, Deisenhofen, Germany). SK-MEL-5 cell line was cultured in MEM (cat. no. 11095-080, Gibco) supplemented with 10% (v/v) heat-inactivated FBS. All cell lines were maintained at 37°C in a humidified incubator with 5% CO_2_. Mycoplasma testing and cell line validation were conducted *via* short tandem repeat profiling by Genetica DNA Laboratories (Burlington, NC, USA) on the following dates: MDA-MB-231 (October 2021), MDA-MB-231-RFP-luc (August 2021), MDA-MB-468 (May 2017), MCF 10A (February 2020), and SK-MEL-5 (February 2022).

All cell lines were confirmed to be mycoplasma-free and authenticated, with a match of 96.43-100% to their reference profiles ^42^.

### 2.3. Human induced pluripotent stem cells culturing conditions

Mono-allelic mEGFP-tagged ACTB WTC human induced pluripotent stem cells (hiPSCs) were obtained from the Coriell Institute, Camden, NJ, USA (cat. no. AICS0016) and cultured in complete mTeSR Plus medium (cat. no. 05825, StemCell Technologies, Vancouver, Canada). The hiPSCs were maintained at 37°C in an atmosphere of 5% CO₂. The cells were confirmed to be free of mycoplasma contamination as of March 2021, verified by Genetica DNA Laboratories (Burlington, NC, USA).

### 2.4. Mini-ring cell viability assay

A total of 8,000 MDA-MB-231 or 10,000 MCF 10A cells were plated per mini-ring in 96-well plates. Mini-rings were formed by pipetting 10 µl per well of single-cell suspensions of cells in a 4:3 ratio of Matrigel (cat. no. 354277, Corning) to complete MammoCult medium (cat. no. 05620, StemCell Technologies) into a ring shape around the rim of the well (cat. no. 92096, TPP, Trasadingen, Switzerland) (**Fig. 2**) as previously described ^43^ ^44^ ^45^. The cell suspensions in the Matrigel:MammoCult mixture were kept on ice throughout the process and vortexed after plating every eight rings to maintain consistency. Plates were incubated at 37°C with 5% CO_2_ for 15 min to allow the rings to solidify. Following solidification, 100 μL of complete MammoCult was added to each well. After a 48-hour incubation period, the medium was replaced with 100 μL of fresh complete MammoCult medium containing either 0.1% DMSO (control) or increasing concentrations (1 nM – 10 μM) of **MON** and its analogs. The medium with DMSO or compounds was refreshed every 24 hours. After 72 hours of treatment, the medium was removed, and wells were washed with 100 μL of pre-warmed DPBS (cat. no. 21-031-CV, Corning). Cells were then released by incubating with 50 μL per well of 5 mg/ml dispase (cat. no. 17105-041, Gibco, Grand Island, NY, USA) for 40 min in 37°C. Subsequently, 10 μL of MTT reagent (5 mg/ml) (cat. no. M2128, Sigma, St. Louis, MO, USA) was added to each well, and the plate was incubated for 24 hours at 37^◦^C with 5% CO_2_. To solubilize the formazan product, 100 μl of 10% SDS in 0.01 M HCl was added to each well, and the plate was incubated for an additional 24 hours at 37^◦^C. Absorbance was measured at 540 nm using BioTek Plate Reader (BioTek Instruments, Inc., Winooski, VT, USA). IC_50_ values were calculated using GraphPad Prism 9 for Windows (GraphPad Software).

### 2.5. Breast cancer organoids acquisition and culturing

MDA-MB-231 BrCa organoids were generated as shown in **Fig. 3A** following our previously published protocol ^19^. Briefly, droplets of a complete MammoCult and Matrigel in 1:4 ratio, containing 10^4^ MDA-MB-231 cells per droplet were generated by pipetting 20 μL of the ice-cold mixture onto a base mold of Parafilm. The droplets were allowed to solidify in a 37°C, 5% CO_2_ humidified incubator for 1 hour, then gently scooped into 24 well plate, with each well containing 1 ml of complete MammoCult (2 droplets per well). The organoids were cultured for 96 hours at 37^◦^C in a 5% CO_2_ environment without agitation. Every other day, 0.5 ml of fresh MammoCult medium was added to each well. After 96 hours, the plate was incubated with rotation at 70 rpm (Orbi-Shaker Jr, Benchmark Scientific, Sayreville, NJ, USA). To prevent necrotic core formation, organoids with an approximate diameter of 3 mm were sectioned into smaller fragments (0.5–1 mm in diameter) after 48 hours using Excelta scissors (cat. no. 17-467-493, 17-456-004, 17-467-497, Fisher Scientific, Waltham, MA, USA) ^46^.

### 2.6. Cell cycle analysis of BrCa organoids

Two BrCa organoids per well were cultured in 24-well plate in 1 ml of complete MammoCult in the presence of 0.1% DMSO (vehicle) or **MON**, or compounds **5**, **6**, **12**, **13**, **15**, **21**, **23** or **32** at concentrations equivalent to 5 x IC_50_ values (**Table 1**) for 24, or 72 hours. Organoids were than harvested using a gentle cell dissociation reagent (cat. no. 07174, StemCell Technologies) containing 0.2% anti-clumping agent, 2-naphtol-6,8-disulfonic acid dipotassium salt (NDA) (cat. no. 439013, Frontier Scientific) and dispersed by gentle pipetting with 1250 µl tip. The cells were subsequently washed with DPBS, fixed with 1 ml of 70% ice-cold ethanol, and stored in-20°C until processing. Cells were pelleted, stained with 300 µl propidium iodide/RNase Staining buffer (cat. no. 550825, BD Biosciences, San Jose, CA, USA), and incubated in the dark at room temperature for 1 hour. DNA content was measured using a FacsAria Ilu Flow Cytometer (BD Biosciences, San Jose, CA, USA), and data were analyzed using FlowJo software.

### 2.7. Wound healing assay

MDA-MB-231 cells were seeded in 60 mm petri dishes (cat. no. 430166, Corning) at a density of 0.3 x 10^6^ per dish and cultured in complete DMEM/F-12 medium for seven days to obtain a confluent cell monolayer. The medium was then aspirated and replaced with serum-free DMEM/F-12. After 24 hours, the medium was removed, and cross-shaped scratches were introduced into the cell monolayer using a 1250 μl pipette tip. The cells were washed twice with DPBS and incubated in medium containing 0.1% DMSO (vehicle), or **MON** (325 nM), compound **5** (42.5 nM), compound **6** (152.5 nM) or compound **12** (65 nM), at concentrations corresponding to half of their respective IC_50_ values (**Table 1**). The wound surface was imaged using the inverted phase contrast microscope EVOS FL Auto Cell Imaging System (Thermo Fisher Scientific) at the beginning of the experiment (0 hours), and after 3, 24, 27, and 48 hours. Three biological replicates were performed. Within each biological replicate, 7 technical replicates per timepoint were averaged together to produce one measurement per timepoint. The resulting 3 measurements per timepoint were used to calculate means ± SDs over time. The wound surface at each time point was normalized to the surface area at 0 hours, which was defined as 100% for each condition, to quantify wound closure.

### 2.8. qRT-PCR analysis

MDA-MB-231 cells (0.1 - 0.2 x 10^6^ cells per well) were seeded in a 6-well plate in complete growth medium and incubated for 48 hours to allow for adhesion. Following medium replacement, the cells were treated with **MON** (325 nM), or analog **5** (45.5 nM), **6** (152.5 nM), or **12** (65 nM) at concentrations equivalent to half of their respective IC_50_ values (**Table 1**). Cells treated with 0.1% DMSO served as the control. After 72 hours of incubation, cells were washed with DPBS, released using 0.05% trypsin, centrifuged, and washed with DPBS. Cell pellets were snap-frozen in liquid nitrogen and stored in-80°C until further processing. Total RNA was isolated using the RNeasy Mini Kit (cat. no. 74104, Qiagen, Hilden, Germany). Genomic DNA removal and reverse transcription were performed using the iScript gDNA Clear cDNA Synthesis Kit (cat. no. 1725035, Bio-Rad, Hercules, CA, USA). The levels of specific transcripts were quantified by qPCR using the following TaqMan assays: SOX2 (Hs01053049_s1), 18s (Hs99999901_s1), and GAPDH (Hs02786624_g1). mRNA transcript levels were normalized to 18s (endogenous control) and calculated using the ΔΔCt method.

### 2.9. Identification of cell surface markers by flow cytometry

MDA-MB-231 cells (0.1 x 10^6^ per well) were seeded in a 6-well plate containing complete growth medium and incubated at 37°C in a 5% CO_2_ humidified incubator for 48 hours to allow for adherence. Afterward, the complete medium was replaced, and the cells were treated with either 0.1% DMSO (vehicle), **MON** (325 nM), or analogs **5** (45.5 nM), **6** (152.5 nM), or **12** (65 nM) at concentrations corresponding to half of their respective IC_50_ values (**Table 1**). After 72 hours of treatment, cells were harvested using enzyme-free cell dissociation buffer (cat. no. 13151014, Gibco) and washed with flow cytometry buffer. The cells were resuspended in flow cytometry buffer and incubated at 1:100 dilution with FITC-conjugated anti-CD24 (cat. no. ab30350, Abcam, Cambridge, Great Britain) and at 1:200 dilution with PE-conjugated anti-CD44 (cat. no. ab46793, Abcam) for 30 min in the dark. After incubation, cells were washed with flow cytometry buffer, resuspended in 300 µl of propidium iodide/RNase Staining buffer, and analyzed using a FacsAria IIIu Flow Cytometer. Data were processed using FlowJo software, with threshold lines set according to isotype controls. The accuracy of the triple immunostaining was confirmed by comparing results with single immunostaining for CD44, CD24, and propidium iodide, respectively.

### 2.10. Clonogenic assay

MDA-MB-231 cells (0.1 x 10^6^ per well) were seeded in 6-well plates (TPP) and incubated for 24 hours at 37°C in a humidified incubator with 5% CO_2_ to allow for adhesion. Following medium replacement, the cells were treated with either 0.1% DMSO (vehicle), or 650 nM **MON**, or 130 nM compound **12** (concentrations equal to their respective IC_50_ values; **Table 1**) for 4, 24, 48, or 72 hours. After treatment, the cells were harvested and reseeded at a density of 250 cells per well (n = 6) in 6-well plates, where they were allowed to proliferate for 21 days. The medium was then removed, and colonies were washed with 1 ml of DPBS, followed by staining with 0.5% crystal violet in methanol for 40 minutes at room temperature on a rocker. Colonies were manually counted, and the results were presented as a percentage relative to the DMSO-treated control cells.

### 2.11. Proteomics analysis

MDA-MB-231 cells (0.9 × 10⁶ per 150 mm petri dish) were treated for 72 hours with 325 nM **MON**, 65 nM compound **12** (corresponding to half of the IC₅₀ values; see **Table 1**), or 0.1% DMSO (control). After treatment, the cells were harvested, washed with DPBS, and resuspended in RIPA buffer supplemented with protease and phosphatase inhibitors. The samples were incubated on ice for 30 minutes, followed by centrifugation at 14,800 rpm for 10 minutes at 4°C. The supernatants were then transferred to a new set of Eppendorf tubes and stored at-80°C until further analysis. The proteomics and phospho-proteomics analysis were performed by the IDeA National Resource for Quantitative Proteomics (https://idearesourceproteomics.org/).

Total protein from each cell lysate was reduced, alkylated, and purified by chloroform/methanol extraction prior to digestion with MS-grade porcine trypsin/LysC (Promega Corporation, Madison, WI, USA). Resulting peptides were labeled using a tandem mass tag 11-plex isobaric label reagent set (Thermo Fisher Scientific, Waltham, MA, USA) and enriched using High-Select TiO_2_ and Fe-NTA phosphopeptide enrichment kits in succession (Thermo) following the manufacturer’s instructions. Both enriched and un-enriched labeled peptides were separated into 46 fractions on a 100 x 1.0 mm Acquity BEH C18 column (Waters, Milford, MA, USA) using an UltiMate 3000 UHPLC system (Thermo) with a 40 min gradient in ratios from 99:1 to 60:40 in buffer A (0.1% formic acid, 0.5% acetonitrile): buffer B (0.1% formic acid, 99.9% acetonitrile) under basic pH conditions, and then consolidated into 18 super-fractions. Both buffers were adjusted to pH 10 with ammonium hydroxide for offline separation.

Each super-fraction was then further separated by reverse phase XSelect CSH C18 2.5 um resin (Waters) on an in-line 150 x 0.075 mm column using an UltiMate 3000 RSLCnano system (Thermo). Peptides were eluted using a 75 min gradient from 98:2 to 60:40 buffer A:B ratio. Eluted peptides were ionized by electrospray (2.4 kV) followed by mass spectrometric analysis on an Orbitrap Eclipse Tribrid mass spectrometer (Thermo) using multi-notch MS3 parameters. MS data were acquired using the FTMS analyzer in top-speed profile mode at a resolution of 120,000 over a range of 375 to 1500 m/z. Following CID activation with normalized collision energy of 31.0, MS/MS data were acquired using the ion trap analyzer in centroid mode and normal mass range. Using synchronous precursor selection, up to 10 MS/MS precursors were selected for HCD activation with normalized collision energy of 55.0, followed by acquisition of MS3 reporter ion data using the FTMS analyzer in profile mode at a resolution of 50,000 over a range of 100-500 m/z.

Proteins were identified and reporter ions quantified by searching the UniprotKB database restricted to Homo sapiens (February 2022) using MaxQuant (Max Planck Institute, version 2.0.1) with a parent ion tolerance of 3 ppm, a fragment ion tolerance of 0.5 Da, a reporter ion tolerance of 0.001 Da, trypsin/P enzyme with 2 missed cleavages, variable modifications including oxidation on M, Acetyl on Protein N-term, and phosphorylation on STY, and fixed modification of Carbamidomethyl on C. Protein identifications were accepted if they could be established with less than 1.0% false discovery. Proteins identified only by modified peptides were removed. Protein probabilities were assigned by the Protein Prophet algorithm^47^. TMT MS3 reporter ion intensity values were analyzed for changes in total protein using the unenriched lysate sample. Phospho(STY) modifications were identified using the samples enriched for phosphorylated peptides. The enriched and un-enriched samples are multiplexed using two TMT10-plex batches, one for the enriched and one for the un-enriched samples.

Following data acquisition and database search, the MS3 reporter ion intensities were normalized using ProteiNorm ^48^. The data was normalized using VSN ^49^ and analyzed using proteoDA to perform statistical analysis using Linear Models for Microarray Data (limma) with empirical Bayes (eBayes) smoothing to the standard errors ^5051^. A similar approach is used for differential analysis of the phosphopeptides, with the addition of a few steps. The phosphosites were filtered to retain only peptides with a localization probability > 75%, filter peptides with zero values, and log2 transformed. Limma was also used for differential analysis. Proteins and phosphopeptides with an FDR-adjusted p-value < 0.05 and an absolute fold change > 2 were considered significant. The mass spectrometry proteomics data have been deposited to the ProteomeXchange Consortium ^52^ via the PRIDE ^53^ partner repository with the dataset identifier PXD060697 and 10.6019/PXD060697.

### 2.12. Immunoblot analysis

MDA-MB-468 cells (0.6 x 10^6^ cells per 100 mm petri dish) were treated for 72 hours with 469 nM **MON**, 332 nM compound **12** (corresponding to half of the IC₅₀ values; data not shown), or 0.1% DMSO (control). SK-MEL-5 cells (0.8 x 10^6^ cells per 100 mm petri dish) were treated for 72 hours with 250 nM **MON**, 103 nM compound **12** (corresponding to half of the IC₅₀ values; data not shown), or 0.1% DMSO (control). After treatment, the cells were washed with DPBS, snap-frozen in liquid nitrogen, and stored at − 80^◦^C until further processing.

Cells were thawed and lysed using lysis buffer containing 25 mM HEPES (pH 7.5), 300 mM NaCl, 0.1% w/v Triton X-100, 1.5 mM MgCl_2_, 0.2 mM EDTA, 0.5 mM DTT, EDTA-free complete protease inhibitor tablets (Roche, Basel, Switzerland), 20 µg/ml aprotinin, 50 µg/ml leupeptin, 10 µM pepstatin, 1 mM phenylmethylsulfonyl fluoride, 20 mM β-glycerophosphate, 1 mM Na_3_VO_4_, and 1 µM okadaic acid. The remaining MDA-MB-231 lysates from the proteomics analysis were used for validation via immunoblotting.

Protein content was evaluated using the Bradford assay, and equal amounts (20 µg) were separated by electrophoresis on 12% (w/v) acrylamide Mini-PROTEAN® precast gels (Bio-Rad). Proteins were electrophoretically transferred onto a PVDF membrane (Immobilon-FL, cat. no. IPFL00010, Merck Millipore, Burlington, MA, USA) and stained with Ponceau S to assess transfer efficiency and verify equal loading. The membrane was then blocked with 5% (w/v) non-fat milk (cat. no. M17200, Research Products International) in Tris-buffered saline containing 0.1% (w/v) TWEEN-20 (TBS-T) for 1 hour at room temperature, followed by incubation for 3 hours at room temperature with primary antibodies against TIMP2 (cat. no. 5738, Cell Signaling Technology, Danvers, MA, USA, 1:1000 dilution) and GAPDH (cat. no. 2118, Cell Signaling Technology, 1:10000 dilution). After washing with 1X TBS-T for five 5-minute intervals, the membrane was incubated with HRP-conjugated goat anti-rabbit IgG (H+L) secondary antibody (cat. no. 170–6515, Bio-Rad, 1:5000 dilution) for overnight at 4^◦^C. Following additional washes with TBS-T, the membrane was exposed to Clarity^TM^ Western ECL Substrate (luminol enhancer solution and peroxide solution, cat. no. 1705060, Bio-Rad) for 5 min. Visualization was performed using the ChemiDoc^TM^ MP Imaging System (Bio-Rad), and quantification were performed using Image J software.

### 2.13. Cerebral organoids acquisition and culturing

Cerebral organoids (COs) were generated using the STEMdiff Cerebral Organoid Kit (cat. no. 08570, StemCell Technologies) following the manufacturer’s instructions and as previously described ^45^. Briefly, when large, compact hiPSCs colonies showing less than 10% differentiation reached 70–80% confluency, they were individually plated in ultra-low attachment 96-well round-bottom plates (cat. no. 7007, Corning) at a density of 9,000 cells per well. Embryoid bodies (EBs) were supplemented every other day with 100 µl of EB Formation Medium per well. On day five, EBs were transferred to 24-well ultra-low attachment plates (cat. no. 3473, Corning) containing 0.5 ml of Induction Medium per well. On day seven, EBs were embedded in cold Matrigel droplets (cat. no. 354277, Corning) and further cultured in 6-well ultra-low attachment plates (12–16 droplets per well) in Expansion Medium (3 ml per well). On day ten, the medium was replaced with Maturation Medium (3 ml per well), and the plates containing organoids were placed on an orbital shaker (Orbi-Shaker Jr, Benchmark Scientific) at 70 rpm in a 37^◦^C, 5% CO_2_ incubator. The medium was replaced every three to four days. On day 51, two representative COs were harvested and fixed with 4% paraformaldehyde for 45 minutes in room temperature, followed by three washes with ice-cold DPBS. The fixed COs were incubated for 1 hour with 15% sucrose at room temperature, after which the solution was replaced with 30% sucrose and incubated overnight at 4^◦^C. The sucrose solution was then removed, and the COs were embedded in Optimal Cutting Temperature Compound (OCT) (cat. no. 4585, Fisher HealthCare, Houston, TX, USA), snap-frozen in liquid nitrogen, and stored at − 20^◦^C until further processing.

### 2.14. Generation and culturing of Hybrid Organoid System: Tumor in Brain Organoid (HOSTBO)

To establish the human Hybrid Organoid System: Tumor in Brain Organoid (HOSTBO), cerebral organoids (hosts) were co-cultured with breast cancer cells (BrCa) stably expressing RFP, modeling the metastatic spread of BrCa to the brain. Individual COs were transferred to non-tissue culture treated 24-well plates (1 CO per well). Each well was supplemented with 2 ml of fresh, complete Maturation Medium containing 10^5^ MDA-MB-231-RFP-luc cells. The COs were incubated for 24 hours at 37^◦^C with 5% CO_2_ without agitation. Following incubation, COs were transferred using wide-bore pipette tips to clean wells in non-tissue culture-treated 24-well plates (one CO per well) and washed twice with 1 ml of DPBS per well. The COs were then cultured in 2 ml of fresh, complete Maturation Medium per well on a 70-rpm orbital shaker at 37^◦^C with 5% CO_2_ for seven days. After seven days, the medium was replaced with 2 ml of fresh, complete Maturation Medium per well, and tumor-bearing COs were cultured for an additional four days in the presence of 0.1% DMSO (control), **MON** (1300 nM), or compound **12** (260 nM), at concentrations corresponding to 2 x their respective IC_50_ values (**Table 1**). Each treatment condition was applied to two individual BrCa-bearing COs. After four days of treatment, the BrCa-bearing COs were harvested and fixed as described in the *Cerebral organoids acquisition and culturing* section.

### 2.15. Histopathological evaluation and immunofluorescence

Immunohistochemical staining was conducted by the UAMS Experimental Pathology Core. For histopathological analysis, OCT-embedded COs were sectioned into 25 µm thick slices and stained with hematoxylin and eosin (H&E) (cat. no. 7231, Richard-Allan Scientific, Thermo Fisher Scientific, Waltham, MA, USA). For immunofluorescence, frozen blocks were cryosectioned into 25 µm thick sections. The sections were permeabilized with 0.2% Triton-X in 1X DPBS for 20 minutes at room temperature in humidified chamber. Primary antibody staining was performed for 2 hours at room temperature in humidified chamber using the following antibodies: Nestin (cat. no. 33475, Cell Signaling Technology, 1:1600), Tubulin β3/TUJ1 (cat. no. 801201, Bio Legend, 1:100), TBR2 (cat. no. 23345, abcam, 1:100), N-cadherin (cat. no. 13116, Cell Signaling Technology, 1:400), SOX2 (cat. no. 4900, Cell Signaling Technology, 1:400), Ki-67 (cat. no. 16667, abcam, 1:250), MAP2 (cat. no. 4542, Cell Signaling Technology, 1:50), cleaved-PARP (cat. no. 5625, Cell Signaling Technology, 1:400), and cleaved-caspase-3 (cat. no. 9664, Cell Signaling Technology, 1:400). The sections were washed three times in DPBS for 5 min at room temperature, followed by incubation with secondary antibodies (cat. no. A21244 (anti-rabbit) or A32728 (anti-mouse), Invitrogen, 1:100) for 1 hour at room temperature in humidified chamber. The sections were mounted onto coverslips using ProLong® Gold antifade reagent with DAPI (cat. no. P36935, Molecular Probes). Images were acquired with a Zeiss LSM880 Confocal System, with Plan-Apochromat 20x/0.8 M27 objective 405 nm, 488 nm, and 633 nm lasers. Analysis was performed using Zeiss Zen 2.6 software. Between 1 and 10 sections were quantified for each treatment condition per CO.

### 2.16. Statistical analysis

GraphPad Prism 9 for Windows (GraphPad Software) was employed for statistical analysis. Welch’s unpaired t-test with Satterthwaite’s degrees-of-freedom correction was used for statistical analysis. All tests employed an unadjusted p<0.05 significance level despite the multiple comparisons, so as not to compromise Type II (false-negative) error in this study. Results are presented as a mean ± SD.

## 3. Conclusions

In the present study, we demonstrated that several esters and urethanes of **MON**, including compounds **5**, **6**, **7**, and **12**, exhibit greater potency against basal-like TNBC cells MDA-MB-231 and higher selectivity for TNBC compared to non-cancerous epithelial cells MCF 10A in a mini-ring organoid-based cell viability assay, compared to parent molecule **MON**. **MON** and selected analogs induced DNA fragmentation in an organoid model of TNBC. Furthermore, **MON** and compounds **6** and **12** significantly inhibited the migration of MDA-MB-231 cells and downregulated the expression of the stemness marker SOX2. **MON** and compound **12** also significantly reduced the CD44^+^/CD24^-/low^ stem-like population of MDA-MB-231 cells and diminished their self-renewal capabilities, as determined by a colony formation assay. Proteomics analysis revealed that these compounds significantly upregulated the expression of TIMP2 in MDA-MB-231 cells, a protein associated with the prevention of metastasis. This effect was consistent across other cancer cell lines, including MDA-MB-468 (a basal-like BrCa subtype) and SK-MEL-5 melanoma cells. Additionally, we developed human HOSTBO to model the metastatic spread of BrCa to the brain. Using this model, we demonstrated that treatment with **MON** and compound **12** led to a significant downregulation of the proliferation marker Ki-67, while compound **12** treatment also resulted in a significant downregulation of SOX2. Taken together, these findings suggest that **MON** and its analogs, particularly compound **12**, exhibit anti-BrCa stem-like cell activity and highlight their potential as therapeutic agents against TNBC and metastatic BrCa.

## Abbreviations

TEA: triethylamine
rt: room temperture
BTC: bis(trichloromethyl) carbonate / triphosgene
DCM: dichloromethane
TBAF: tetra-n-butylammonium fluoride
THF: tetrhydrofurane
anh.: anhydrous
aq.: aqueous
sol.: solution
DMAP: 4-Dimethylaminopyridine
PPh3: triphenylphosphine
DCC: N,N′-Dicyclohexylcarbodiimide
Py: *Pyridine*
p-TSA: *p-Toluenesulfonic acid*
DBU: *1,8-Diazabicyclo[5.4.0]undec-7-ene*
HOBt: Hydroxybenzotriazole
DMF: dimethylformamide.

## Acknowledgements

This work was supported by the UAMS Translational Research Institute through grants KL2 TR003108, K12 TR004924, UM1 TR004909, and UL1 TR003107 from the NIH National Center for Advancing Translational Sciences; Barton Pilot grant (to AU); the Arkansas Breast Cancer Research Program (to AU and AMM); R01 HD093461 and RO1 DK139476 (to AMM); and R24 GM137786. We thank Ms. Andrea Harris from the UAMS Flow Cytometry Core, Ms. Jennifer James from the UAMS Experimental Pathology Core, and Mr. Jeff Kamykowski from the UAMS Digital Microscopy Core Laboratory. We thank Drew Seale, a summer student, for helping with the immunofluorescence staining of cerebral organoids. We would also like to thank Drs. Eric Norris and HoWon Kim from StemCell Technologies for valuable discussions regarding iPSC culture and CO generation, and Dr. Amanda Linkous for her consultations and helpful suggestions for the development of the GLICO model. Reactome Pathway Database (reactome.org) was used to identify significantly differentially expressed pathways in proteomics and phosphoproteomics experiments. Some elements of the figures were created using BioRender (https://www.biorender.com/).

## Author contributions

AU performed the experiments, acquired funding, analyzed and visualized the data, and wrote the first draft of the manuscript. BH sectioned cerebral organoids. ES supported statistical analysis and revised the manuscript. MRR contributed to figure creation and pathway analysis. JSN provided pathology analysis, captured H&E sections, and revised the manuscript. EUY contributed to pathology analysis. MJ synthesized and purified compounds. GK synthesized and purified compounds, prepared figures, and contributed to manuscript writing. NS synthesized and purified the compounds. AH supervised compound synthesis, purification, and identification, and supported funding acquisition. MBN provided consultation. TCC participated in funding acquisition and edited the manuscript. SP provided consultation. RLE supported funding acquisition. MCM supported funding acquisition. AKT provided consultation. TK provided consultation, supported data interpretation, and edited the manuscript. AJT supported funding acquisition, supervised the project, and contributed to data interpretation. AMM acquired funding, participated in experimental design, contributed to data analysis and interpretation, and supervised the project.

## Supporting materials

Additional experimental data including mass spectrometry proteomics data and H&E and immunofluorescence images of cerebral organoids.

