## Supplementary Material for "Monensin and Its Analogs Exhibit Activity Against Breast Cancer Stem-Like Cells in an Organoid Model"

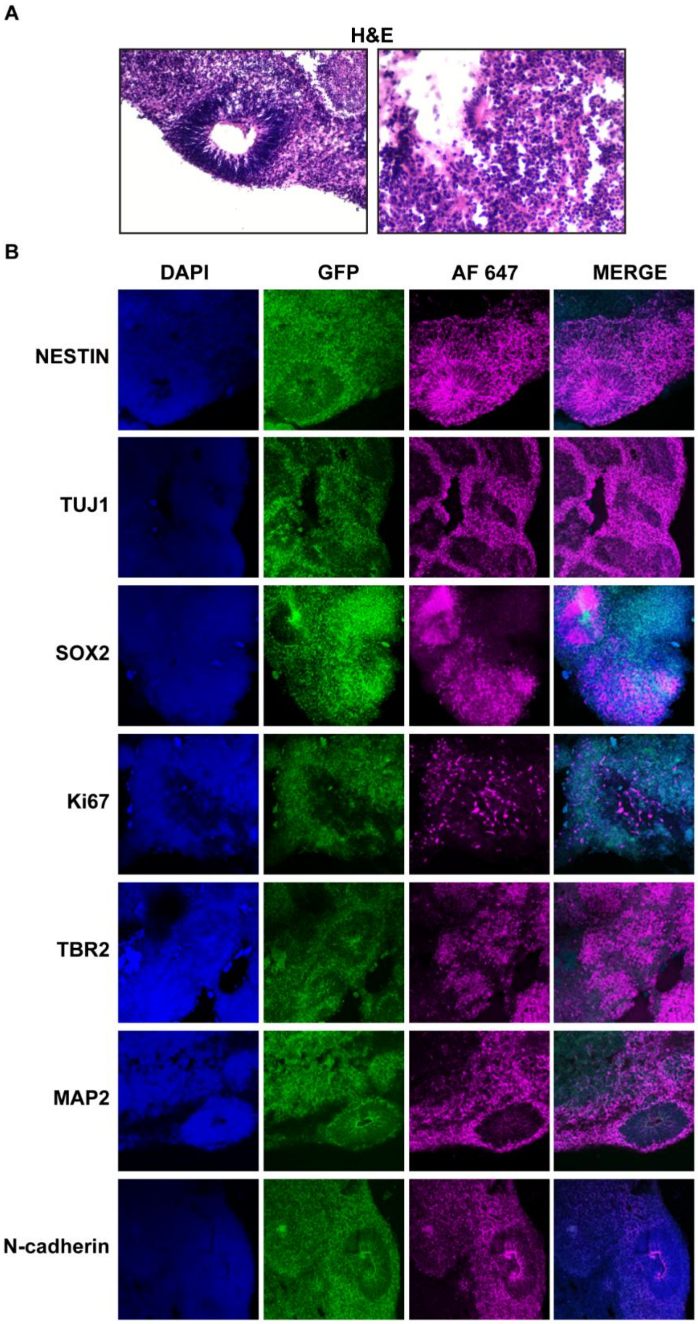


**Figure S1**. Cerebral organoids validation. **A**. H&E staining of normal cerebral organoids at 51 days of development, captured at 20X magnification; **B**. Immunofluorescence staining showing DAPI (nuclei), GFP, and Alexa Fluor 647 (AF 647), a secondary antibody used for markers including Nestin, Tuj1, Sox2, Ki-67, TBR2, MAP2, and N-cadherin, captured at 20X magnification.
